# OrganoidChip+: a Microfluidic Platform for Culturing, Staining, Immobilization, and High-Content Imaging of Adult Stem Cell-Derived Organoids

**DOI:** 10.1101/2025.06.02.657499

**Authors:** Khashayar Moshksayan, Rishika Khanna, Berk Camli, Zeynep Nevra Yakay, Anirudha Harihara, Christopher Zdyrski, Sudip Mondal, Evan Hegarty, Jonathan P. Mochel, Karin Allenspach, Adela Ben-Yakar

**Affiliations:** Department of Mechanical Engineering, University of Texas at Austin, Austin, TX; Department of Biomedical Engineering, University of Texas at Austin, Austin, TX; Department of Pathology, University of Georgia, Athens, GA; Precision One Health Initiative, University of Georgia, Athens, GA; Department of Electrical and Computer Engineering, University of Texas at Austin, Austin, TX

## Abstract

High-content imaging (HCI) and analysis are the keys for advancing our understanding of the science behind organogenesis. To this end, culturing adult stem cell-derived organoids (ASOs) in a platform that also enables live imaging, staining, immobilization, and fast high-resolution imaging is crucial. However, existing platforms only partially satisfy these requirements. In this study, we present the OrganoidChip+, an all-in-one microfluidic device designed to integrate both culturing and HCI of ASOs all within one platform. We previously developed the OrganoidChip as a robust imaging tool. Now, the OrganoidChip+ incorporates several additional features for culturing organoids in addition to fluorescence staining and imaging without the need for sample transfer. The organoids grown within a culture chamber are stained and then transferred to immobilization chambers for blur-free, high-resolution imaging at predetermined locations. We cultured adult stem cell-derived intestinal organoids in the chip for 7 days and tracked growth rates of each organoid using intermittent brightfield images, followed by multiple image-based assays, including viability assay using widefield fluorescence imaging, a redox ratio assay using label-free, two-color, two-photon microscopy, and immunofluorescence assays using confocal microscopy. These assays serve as proof-of-concept to showcase the chip’s capabilities in HCI of ASOs. Organoids cultured in the chip exhibited superior average growth rates over those in traditional Matrigel dome cultures, off-chip. Viability and redox ratio measurements of on-chip organoids were comparable or slightly better than their off-chip counterparts. Confocal imaging further confirmed that the OrganoidChip+ supports robust organoid culture while enabling detailed, high-resolution analysis. This all-in-one platform holds great potential for advancing ASO-based research, offering a scalable and cost-effective solution for HCI and analysis in organogenesis, drug screening, and disease modeling.

## Introduction

Organoids are miniaturized three-dimensional (3D) structures that mimic the characteristics of tissues. To study organoids’ development, cellular heterogeneity, and drug response, it is necessary to use platforms that enable culturing, staining, and fast High-Content Imaging (HCI), within the same platform. While HCI is a common practice for *in vitro* studies using 2D culture models, the currently available technologies only partially meet the requirements for HCI of 3D adult stem cell-derived organoids (ASOs). One important challenge with the conventional Matrigel dome cultures is the organoids’ random distribution that places them far from the imaging substrate. To capture all the organoids, users will need to image the entire dome volume using a large number of field of views (FOVs) during time-consuming sessions. Furthermore, when seeding ASOs in multiwell plates, contact with the well walls often causes organoids to reside at the edges, leading to aberrations and non-uniform organoid distribution in the well. Both problems necessitate the transfer of organoids to a secondary platform with a thin substrate for high-resolution imaging. Transferring organoids incurs several disadvantages, including altering their morphology due to cell shedding in un-healthy organoids, losing track of specific organoids during the transition from the primary to the secondary platform, agglomeration of multiple ASOs into individual clumps, and the loss of organoids as they adhere to liquid handling tools. Another challenge is that when released from the hydrogel-based extracellular matrix (ECM), organoids are not fully immobilized for performing imaging-based assays, e.g. immunofluorescence. Organoids that are not fully immobilized can drift during fast imaging sessions for high-throughput screening, due to the movements of the microscope stage or the reagents in the imaging wells which result in blurry images.

Numerous novel devices have been proposed for on-platform culturing and imaging of ASOs, yet they remain subject to varying degrees of the previously mentioned limitations. While these devices provide more control over spatial distribution of ASOs, many lack a thin substrate^1,2^ or require complex systems for high-resolution imaging^2–4^. For example, the OCTOPUS platform^4^ features a device with height of 1 mm, which necessitates capturing of a large number of z-images when using high numerical aperture (NA) objectives. Generally, air objectives with high NA (≥ 0.75) have a limited working distance of ∼ 0.6 mm. As a result, water immersion objectives become necessary, adding operational complexity and expense for the automated water injection systems. To address this challenge, Schuster *et al.* developed a microfluidic device with a restricted culture chamber height of 610 µm^1^. Their approach eliminated the need to transfer organoids to a secondary imaging device. However, high-resolution imaging remained impractical due to lack of a thin substrate, a limitation imposed by its specialized fabrication method.

Additional devices, such as the OrganoPlate^5^ (MIMETAS) and the idenTx^6^ (AIM Biotech) are promising solutions for disease modeling, staining, and high-resolution imaging of ASOs with long-term culture capabilities. With simple design and fabrication, these platforms address most of the mentioned barriers, however, they cannot provide immobilization that is required for HCI of ASOs upon digestion of the hydrogel. Applications such as immunofluorescence staining necessitate the digestion of hydrogel for better reagent penetration and for avoiding non-specific binding. Recently, the JeWell system was able to address the immobilization challenge and facilitate HCI of organoids^7^. The authors implemented embryoid body formation and differentiation to generate organoids in the semi-pyramidal gold coated Jewells that enables light-sheet microscopy. The device provides full immobilization of organoids upon Matrigel digestion and allows for on-platform immunofluorescence staining. Despite all the advantages, the authors did not show brightfield imaging using this device, most likely due to the pyramidal structure blocking transmitted light. Furthermore, this platform requires a customized light-sheet microscope and fluorescently labelled cells for imaging, which may limit accessibility and broader adoption. The ability to perform brightfield imaging is crucial in organoid screening as it facilitates monitoring of organoid growth rate. Furthermore, machine learning algorithms have proven brightfield imaging to be a valuable tool by successfully assessing organoid viability after drug treatment using brightfield images.

To address the challenges mentioned, we aim to fulfill six key pillars for developing an effective culturing and imaging platform that enables high-throughput and HCI of ASOs: (1) transferless culturing, immunofluorescence staining, and high-resolution imaging; (2) limited span of organoid location in the z-direction; (3) pre-determined organoid locations to enable high-throughput assays; (4) complete organoid immobilization; (5) a flat, thin substrate for high-resolution imaging; and (6) a scalable and cost effective design for high-throughput screening. To this end, we leveraged the capabilities of our previously designed platform, the OrganoidChip^8,9^, in the new design, the OrganoidChip+, to fulfill the six pillars of HCI in a series of proof-of-concept experiments using canine-derived colon ASOs. Previously, the OrganoidChip served as an imaging tool with trapping areas (TAs) designed for organoid immobilization and high-resolution imaging. In this study, we enhanced the chip’s functionality by enabling on-chip culturing of organoids. The modified chip design includes a perfusion channel on each side of the culture chamber to ensure a more uniform delivery of reagents to the cultured organoids.

The average growth rates of organoids cultured on-chip were comparable to those cultured off-chip in multiwell plates, indicating a healthy culture environment for the ASOs within the device. Following culturing of organoids, we used high-resolution imaging to implement three image-based assays including organoid viability assay, redox ratio assessment using label-free, two-color, two-photon microscopy, and immunofluorescence assay using confocal microscopy, all on the chip. These studies allowed us to demonstrate high cellular viability and redox ratios as well as relevant cellular phenotypes by enabling culturing and transferless staining and high-resolution imaging. We anticipate that OrganoidChip+ will play a pivotal role in ASO research by offering real-time monitoring of organoids and performing high-content endpoint analysis of individual organoids, all in one platform.

## Results and discussion

### Device design and operation

The OrganoidChip+ consists of an inlet, a culture chamber, two side perfusion channels, six trapping areas (TAs), and a serpentine exit channel to create hydrodynamic resistance (**Fig. 1a**). The height of the culture chamber is restricted to 550 µm to facilitate organoid growth close to the thin glass substrate, enabling high-resolution imaging with high-NA objectives. As shown in **Figs. 1b** and **1c**, the intestinal cell-Matrigel suspension is seeded into the culture chamber by carefully inserting a bent Luer stub into the culture chamber through the inlet and dispensing ∼ 5 µL of the suspension. The culture chamber is separated from the perfusion channels by two pendent walls that leave 100 µm gap underneath to deliver the reagents to the culture chamber. The pendant walls also prevent the liquid Matrigel suspension from entering and clogging the perfusion channels during cell seeding. The growing organoids in the culture chamber are exposed to reagents (nutrients, dyes, etc.) from all directions including the inlet (top), the perfusion channels (sides), and the TAs (bottom) (**Fig. 1b**). There are two glass barrels, each with 230 µL volume, one mounted at the inlet, to serve as the nutrient and reagent reservoir for organoids, and one at the outlet of the chip (**Fig. 1c** and **Supplementary Fig. S1**).

**Figure 1:**
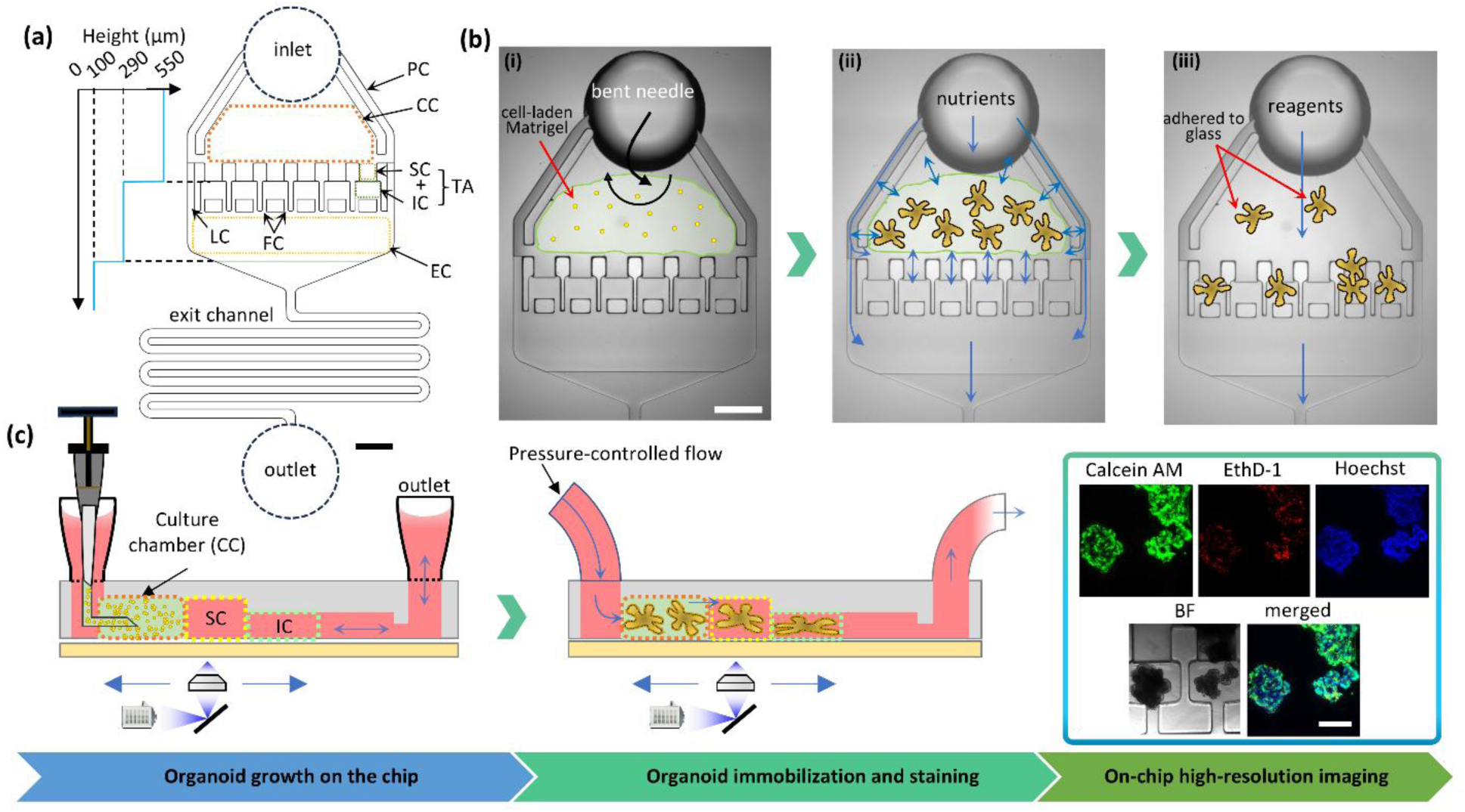
OrganoidChip+’s dimensions and principle of work. **(a)** The chip schematic depicting its height dimensions and various compartments such as the inlet, perfusion channel (PC), culture chamber (CC), trapping areas (TAs) consisting of staging chambers (SCs) and immobilization chambers (ICs), filter channels (FCs), exit chamber (EC), and serpentine exit channel. The serpentine exit channel with a length of 43.5 mm and a width of 215 µm generates a hydrodynamic resistance of 3 × 10^12^ (N.s/m^5^) to avoid high flow rates and shear stress inside the chip. **(b)** Various steps for cell seeding (i), organoid culture (ii), and immobilization (iii) in the chip. (i) For seeding, the Luer stub is inserted into the culture chamber, the cell suspension is dispensed with a rotating movement to fill the entire culture chamber. (ii) Organoids grow efficiently while accessing nutrients from all direction surrounding the culture chamber. (iii) Matrigel is digested, and organoids are pushed into the TAs for immobilization while some organoids are natively immobilized by adhering to the glass in the culture chamber. **(c)** Side view of the same steps depicted in (b). The organoids in the culture chamber are imaged using brightfield microscopy every day to track their growth. After 7 days of culturing, Matrigel is digested to enable organoid immobilization within TAs. After immobilization, organoids can be fluorescently labelled and imaged at high-resolution on the chip. Scale bars represent 1 mm in (a-b) and 400 µm in (c).

Organoids were grown for 7 days on the chip before performing viability assays, redox ratio imaging, and immunofluorescence staining. For the viability and immunofluorescence staining assays, we first digested the Matrigel, proceeded with staining steps, and then moved the freed organoids into the TAs using a flow-induced drag as described previously^8^ (**Videos S1 & S2**, see also Methods). Some organoids adhered to the glass during their growth and, therefore, remained attached to the glass during viability staining and immobilization process. These organoids were naturally immobilized in their native locations without requiring further immobilization (**Fig. 1b**). On the other hand, most organoids that were fixed with 4% Paraformaldehyde (PFA) prior to staining lost their adhesion. PFA fixation, which crosslinks the proteins by the Aldehyde groups^10^, significantly reduces their surface stickiness, allowing them to detach from their native location and move to the TAs during immobilization (**Video S1**). Redox ratio imaging was performed on organoids still embedded in Matrigel on Day 7 post-seeding.

The TAs, consisting of staging and immobilization chambers, enable full immobilization of the organoids and provide predetermined locations for fast high-resolution imaging (**Fig. 1a**). The immobilization chambers have a reduced height of 290 µm, which allows for immobilization of smaller organoids and further reduces the number of required z-images compared to the staging chambers. The organoids within the immobilization chambers are immobilized by friction forces, as they are gently compressed between the ceiling and the bottom glass (**Fig. 1c**).

To prevent pressure build-up within the chip when all TAs are filled with organoids, we implemented two precautionary actions. First, we adjust the upstream pressure to precisely control the flow rate in the fluidic setup. Thus, when TAs are filled and hydraulic resistance increases, the flow rate decreases to avoid high shear stress on organoids. Second, we designed lateral channels that bypass the TAs by connecting the culture chamber to the exit chamber. These two channels provide an alternate flow path when the TAs are fully occupied by organoids. Importantly, lateral channels remain free of clogging since the device’s flow dynamics and perfusion walls guide the organoids preferentially toward the TAs.

The filter channels connect the immobilization chambers to the exit chamber while preventing organoid escape. We developed two immobilization chamber designs that differed in the number of filter channels: one with a single channel and one with two channels (**Supplementary Fig. S1**). Organoids trapped in the immobilization chambers tend to localize near the inlet of a filter channel, imposing flow-induced shear stress on the peripheral cells. The motivation for having two filter channels was to reduce this shear stress by allowing some flow to bypass the organoid through the unoccupied filter channel.

However, incorporating the lateral channels into the current OrganoidChip design alleviated the need for having two filter channels. This is because a large portion of the flow freely passes through the lateral channels, therefore, decreasing the shear stress on organoids that are trapped in immobilization chambers with a single filter channel. Since both designs (**Fig. 1a** and **Supplementary Fig. S1**) exhibited comparable performance, they were used interchangeably throughput the study.

### Comparison of organoid growth between off- and on-chip cultures

We observed that single organoids can grow from either a single cell or an agglomeration of two or more cells when suspended in Matrigel at seeding. We refer to these single cells or agglomerations as clumps when discussing the seeding density. The seeding density was optimized to yield an average of 8 to 12 organoids per chip that are larger than 400 µm in size, while maintaining growth rates similar to those of off-chip counterparts (see **Supplementary Fig. S2** for details). We used an optimized seeding density of 1.5 × 10^5^ clumps/mL of Matrigel and refreshed the culture medium in both the conventional culture wells and the on-chip reservoirs every 48 hours to maintain consistent culture conditions across all experiments. Brightfield imaging was performed over 7 days to track organoid growth (**Fig. 2a**). The projected area of each organoid was determined from binarized brightfield images (**Fig. 2b** and **Supplementary Fig. S3**). The daily growth rate was defined as the ratio of the organoid’s projected area on a given day to the area of the initial clump at seeding on Day 0 (**Fig. 2c** and **Supplementary Eq. S1**). This definition accounted for the variability in clump sizes at seeding. The daily growth rates, averaged over 7 experiments for both off-chip and on-chip groups, are presented in **Fig. 2d** (**Supplementary Eqs. S1-S3**). Statistical analysis indicates no significant difference between the two conditions, except for Day 7, where higher average growth rates were observed on-chip.

**Figure 2:**
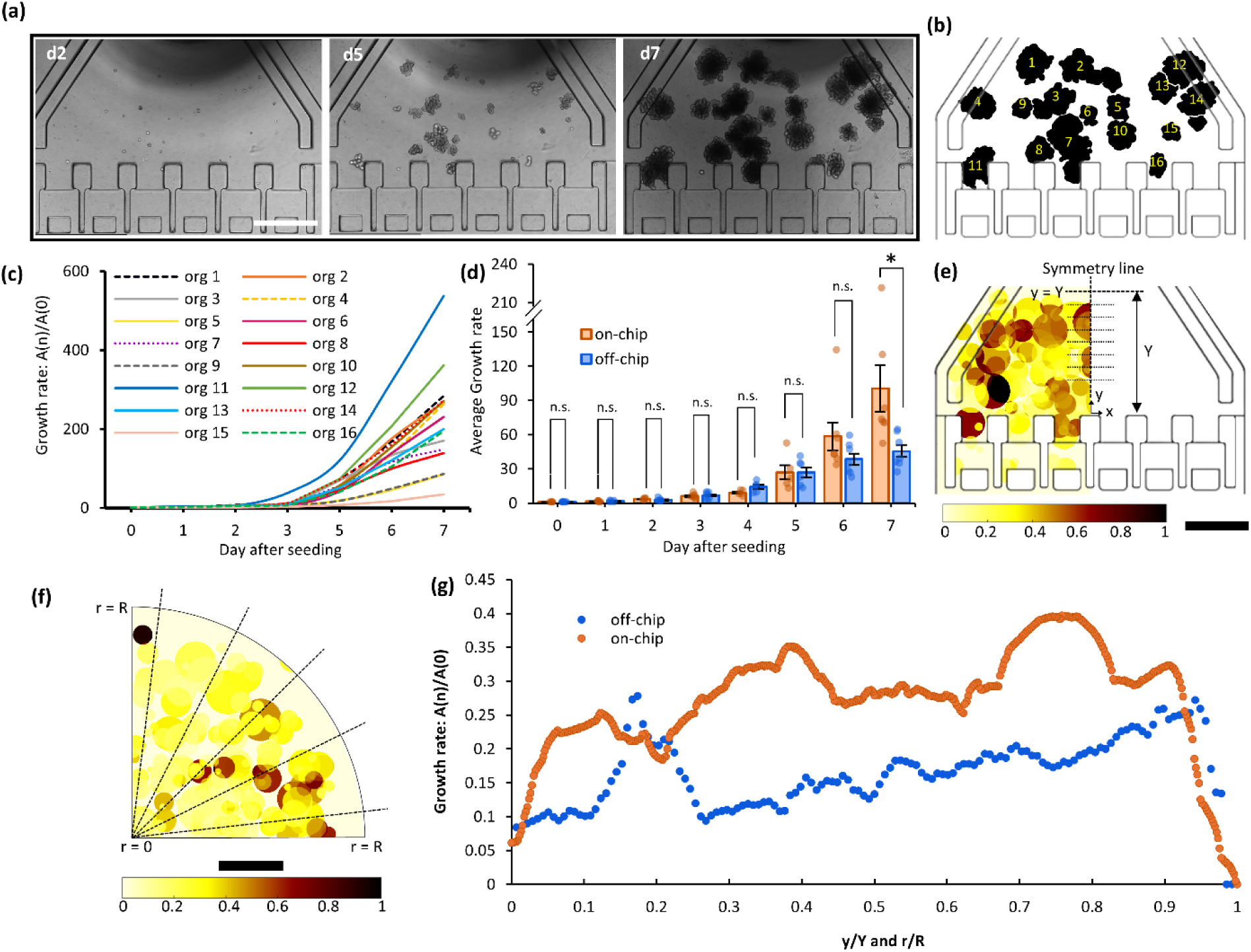
The growth rate of organoids cultured off- and on-chip. **(a)** Brightfield images on Days 2 (d2), 5 (d5), and 7 (d7) after seeding. **(b)** Segmented binary image corresponding to Day 7 of (a). **(c)** Organoids’ areas as calculated from segmented images in (b) and normalized by their Day 0 values. **(d)** Organoid growth rates between Days 0 to 7 for the off- and on-chip experiments. Each dot represents the mean growth rate of all organoids in a single experiment and the bars depict the average growth rates across 7 experiments. Each experiment consists of one chip and at least one quarter of a Matrigel dome. Error bars represent standard deviations. Heatmaps **(e)** and **(f)** show the organoids as circles with their equivalent radius for 7 experiments in the culture chamber and the dome, respectively. Circles are color-coded to depict the normalized growth rates. **(g)** Heatmap values on the dotted lines in (e) and (f) were averaged and plotted along y-direction in (e) and radially in (f). Not all lines are displayed in (f), as a total of 91 lines were used, with angles relative to the horizontal edge of the quarter ranging from 0° to 90° in 1- degree increments. Scale bars in (a) and (e-f) represent 1 mm.

Next, we studied organoid growth rates in both Matrigel domes and chips, specifically on Day 7. **Figures 2e-f** show two heatmaps that depict the organoid growth rates on Day 7 on-chip and in the 25 µL Matrigel dome cultures, respectively, averaged across 7 experiments (see **Supplementary Eqs. S4-S5** for details). The heatmaps are presented on similar dimensional and color scales to help compare the size and growth rates of the two cultures. To do so, the heatmap values were normalized with respect to the maximum growth rate value found between the two conditions, which was observed on-chip. The on-chip results reveal growth rates that are higher in magnitude than the dome cultures (**Fig. 2g**). In dome cultures, the growth rate varies radially, significantly decreasing between the periphery and the central regions with the periphery exhibiting faster growth^4,11^. This variation can be attributed to the limited diffusion of nutrients and metabolites through the Matrigel matrix, which creates a gradient favoring regions closer to the nutrient source. The reduction in the growth rates is also seen as moving away from the inlet reservoir in the chip, albeit being less than that of the domes. This is due to the smaller dimensions of the culture chamber and the availability of nutrients to the organoids from all directions by the perfusion channels and the TAs that bring the culture medium around the culture chamber. The critical role of TAs in nutrient delivery is validated by the observed growth rate values of organoids grown in the bottom-central region of the culture chamber, near the TAs, which is located far from the perfusion channels and the inlet reservoir. In that area, there are organoids with high growth rates (orange) that were fed with the available nutrients diffusing into the Matrigel from the culture media in the TAs. Overall, growth rates in the chip are higher than the dome culture which agrees with the daily growth rate results in **Fig. 2d**.

### OrganoidChip+ does not affect the viability of organoids cultured on-chip

Fluorescence viability assays are standard methods for evaluating cellular health and functionality *in vitro*. To compare the viability of 7-day-old organoids grown off- and on-chip, we stained them with Calcein AM, EthD-1, and Hoechst. We first digested the Matrigel in the chip and then immobilized the liberated organoids in the TAs using our fluidic setup (see Methods). Some on-chip organoids tend to adhere to the glass substrate during growth, staying immobilized at their native location. The remaining organoids, although showing adherent properties, can be gently pushed into the TAs and immobilized in the staging or immobilization chambers for high-resolution imaging (**Figs. 3a-b**). **Figure 3a** shows the organoids in their native culture before Matrigel digestion, while **Fig. 3b** shows the same chip after Matrigel digestion and immobilization, with some organoids entering the TAs.

**Figure 3:**
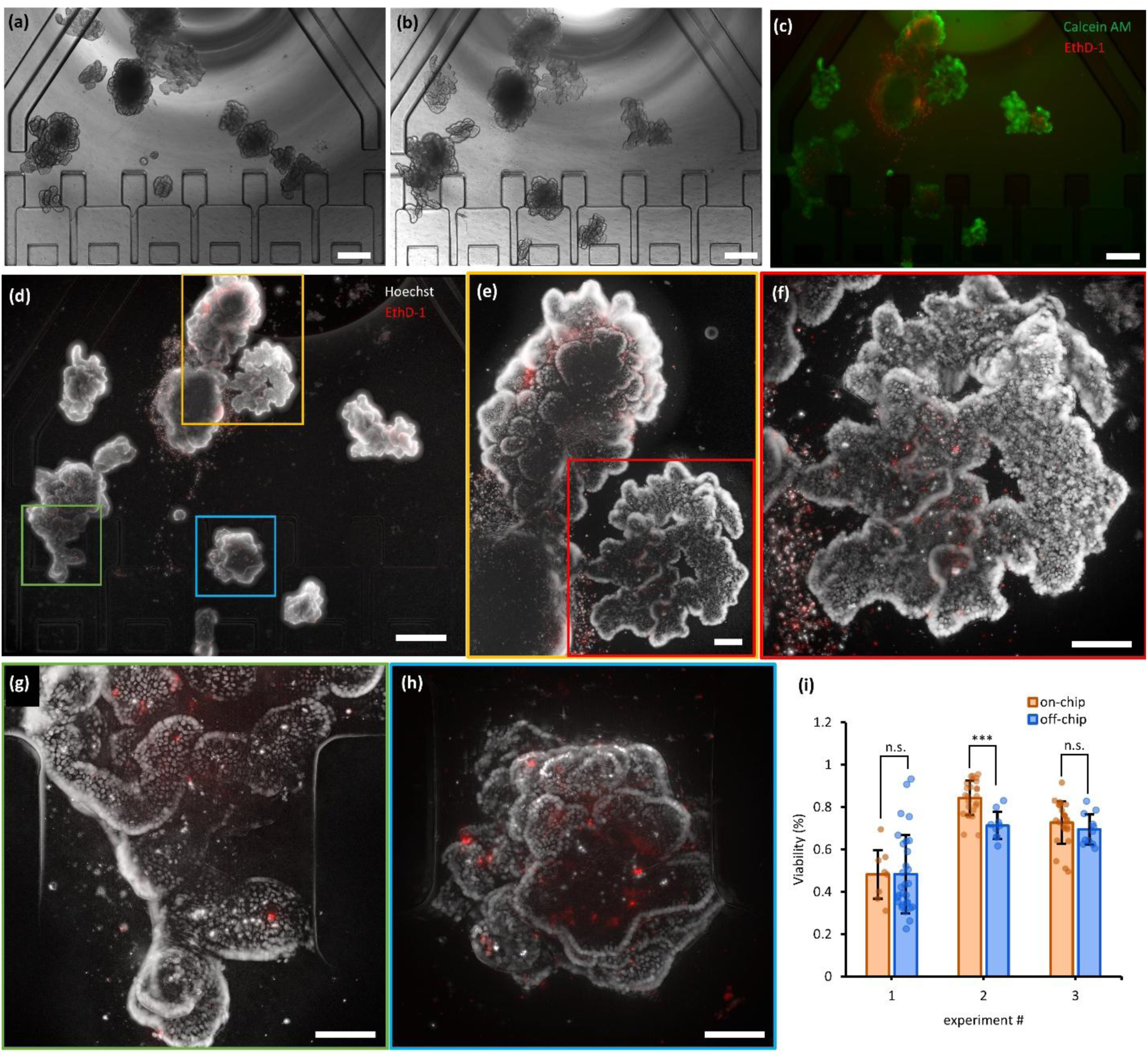
Widefield imaging of fluorescently labelled organoids grown in the OrganoidChip+. (a-b) Images of on-chip organoids on Day 7, **(a)** before digesting the Matrigel and **(b)** after immobilization. **(c)** Images of organoids that are stained with Calcein AM (green) and EthD-1 (red) and imaged with a 0.16 NA, 4× objective. **(d-h)** Images of organoids that are cultured and stained on the same chip with Hoechst (gray) and EthD-1 (red) to show nuclei of all cells and dead cells, respectively. Image in (d) corresponds to the same chip shown in (a-b), indicating the FOVs imaged using higher NA objectives in (e-h). Image in (e) is captured with 0.3 NA, 10× objective, and those in (f-h) are captured using 0.75 NA, 20× objective. The high-resolution imaging was able to resolve single nuclei of the live and dead cells in organoids. **(i)** Depicts viability of organoids across three experiments. Each experiment consists of two chips and two wells. Each dot represents the viability of one organoid. The viability values are not statistically different except for the second experiment (two-way *t*-test). The error bars show standard deviations. Scale bars represent 400 µm in (a-d) and 100 µm in (e-h).

Next, we performed three rounds of fluorescence widefield imaging at multiple z-planes using three different objectives. We began with a low resolution 0.16 NA, 4× objective to find the location of each organoid after immobilization (**Fig. 3d** and **Supplementary Fig. S4**). We then switched to a 0.3 NA, 10× (**Fig. 3e**) and a high-resolution 0.75 NA, 20× objective (**Figs. 3f-h**). High-resolution imaging was able to resolve the nuclei of live and dead cells stained with Hoechst and EthD-1, respectively, and captured the organoids’ morphology (**Figs. 3f-h**). The ability to resolve the nuclei is crucial since it enables counting the number of fluorescently labelled live and dead cells, providing a more accurate assessment of cell viability compared to indirect methods. For example, the CellTiter-Glo® assay indirectly estimates viability by measuring luminescence intensity that reflects the amount of ATP present in the culture, rather than assessing cell numbers directly^12^. Furthermore, *in situ* high-resolution imaging allows the detection of the fluorescently labelled target phenotypes without the need for organoid dissociation for downstream analysis techniques such as FACS. **Figures 3f-h** show healthy intestinal crypt-like structures with a few EthD-1^+^ cells immobilized in both the culture chamber and the TAs.

The statistics for three separate experiments are represented in **Fig. 3i**, where each bar shows the average viability for all organoids grown in the off- and on-chip experimental groups (**Eq. 1**). The results indicate that the on-chip organoid viability is either equal to or higher than that of the off-chip groups. The slightly lower off-chip viability, especially in the second experiment, may be attributed to the transfer of organoids from their native culture platform to an imaging platform (e.g. an 8-well µ-slide) for staining and imaging. This transfer process involves multiple pipetting steps, which can lead to cell shedding, particularly in necrotic organoids, and may cause alterations in organoid morphology^8^.

### Redox ratio imaging reveals consistent off- and on-chip levels

NADH and FAD play crucial roles in epithelial cell growth, metabolic activity, and apoptosis through their involvement in energy metabolism and oxidation-reduction reactions. During cell growth, NADH is essential for ATP generation via the electron transport chain, while FAD participates in oxidative metabolism and mitochondrial redox reactions critical for cellular development^13^. Studies show that changes in NADH and FAD levels can be linked to mitochondrial function impairment and oxidative stress that potentially trigger cell death pathways^14^. The ratio of the NADH and FAD signal intensity (NADH/FAD), known as the redox ratio, serves as an index of mitochondrial metabolic state in cancer tissues^15^, with its value declining following treatments with anticancer drugs such as Doxorubicin (Dox), as verified via redox ratio imaging^16^. Dox has been shown to elevate oxidative stress in cancerous tissues^17^ and colon organoids and to trigger apoptotic pathways as verified by caspase 3/7 activation assay, elsewhere^18^. To date, no published studies have reported on redox ratio imaging in healthy intestinal organoids treated with or without Dox. We therefore hypothesized that, because the pathways leading to reactive oxygen species production and apoptosis after Dox exposure are shared between healthy intestinal organoids and cancerous tissues, a similar decrease in redox ratio would occur. To examine this hypothesis, we evaluated the chip’s capability for investigating the redox ratios in intestinal organoids cultured both off- and on-chip, with and without Dox treatment, using label-free, two-color, two-photon microscopy.

Organoids were treated with vehicle control (DMSO, 0.1% v/v) and 0.3 and 2 µM concentrations of Dox five days after seeding for 48 hours both off- and on-chip to induce dose-dependent mitochondrial damage and apoptosis. Images were captured using an in-house-built two-photon microscope equipped with a high-resolution 0.75 NA, 20× (Nikon Olan Apo) objective and coupled with a Ti:sapphire laser (Mai Tai, Spectra-Physics) to excite the NADH and FAD molecules^19^. We performed image analyses and calculated the redox ratios as described in **Supplementary Fig. S5** and **Eqs. S6-S12**. The redox ratio value for each organoid was normalized to the pooled average redox ratio value of the off- and on-chip vehicle control groups of the same experiment to compensate for the batch-to-batch variation of the samples between experiments^20^ (**Supplementary Eqs. S11-S12**).

Figure 4a shows the normalized redox ratio heatmaps for both the vehicle control and Dox-treated organoids. The high-resolution images highlight various organoid-specific architectural compartments such as the intestinal crypts and lumens. The redox ratio qualitatively decreases at higher Dox concentrations due to the higher apoptotic levels in those organoids, indicating a shift towards a more oxidized state^21^, compared to healthy organoids that are in a reduced state with high redox ratios^15^. For instance, the on-chip organoid treated with 2 µM Dox shows high FAD and low NADH signals from apoptotic cells that were shed into the lumen, which resulted in low redox ratio values overall and in that region specifically. Furthermore, the off-chip organoid treated with 2 µM Dox in Fig. 4a lost the integrity of the epithelial lining due to high cell apoptosis, as confirmed by other studies^22,23^. Figure 4b depicts the morphology of the intestinal lumen (orange arrows) and the epithelial cell lining architecture (white arrows) that were changed in the organoids treated with 0.3 µM compared to the vehicle control. We also observed that the organoid size as well as the number of crypts and lumens decrease at higher concentrations, which likely stems from the negative effects of Dox treatment on stem cell function and proliferation. **Videos S3-S5** show images captured at the two wavelengths (745 nm and 860 nm) as traveling deeper into the vehicle control organoids and those treated with 0.3 µM and 2 µM Dox concentrations on the chip.

**Figure 4:**
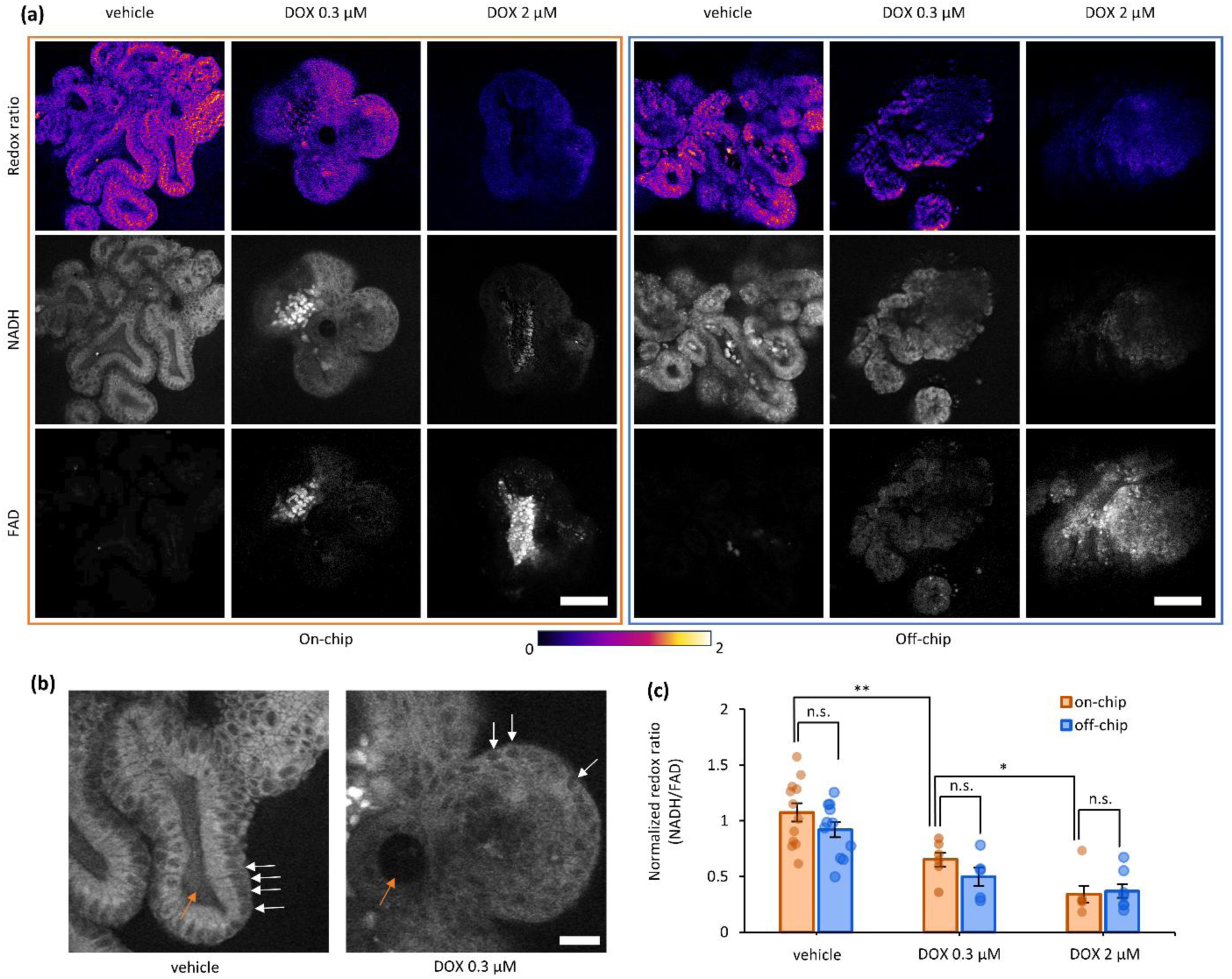
Redox ratio imaging using label-free, two-color, two-photon microscopy. Three organoid groups were treated with 0.1% DMSO (vehicle control), and 0.3 µM and 2 µM Dox concentrations on Day 5 after seeding and incubated for 48 hours prior to imaging. Each treatment condition was conducted both off-chip and on-chip. **(a)** Sample images from off- and on-chip organoids showing the NADH, FAD, and redox ratio (NADH/FAD) extracted from the autofluorescence images collected at the excitation wavelengths of 745 nm (exciting both NADH and FAD) and 860 nm (exciting only FAD). **(b)** Magnified view of NADH images captured from on-chip vehicle control and 0.3 µM Dox-treated organoids. Images show the organoid lumens with orange arrows and epithelial cell linings with white arrows. **(c)** Shows the normalized redox ratio values where each data point corresponds to an organoid and bars denote the mean values. Each Dox-treated group was repeated three times with the corresponding vehicle DMSO control (n = 2 organoids per group). Error bars represent standard error of the mean (SEM). Scare bars represent 100 µm in (a) and 30 µm in (b).

Figure 4c shows the normalized redox ratio values which decrease by increasing Dox concentrations, confirming the results in Fig. 4a. Various experimental groups were compared for the equality of the variances and means using *F*-test and *t*-test (see **Supplementary Tables S3-S4** for the *p*-values). There was no statistically significant difference between the off- and on-chip data for the same Dox concentrations. The mean values were also compared across various concentrations within the off- and on-chip groups. All differences were found to be statistically significant, except between the off-chip 0.3 µM and 2 µM concentration groups. This exception likely originates from the additional preparation steps such as pipetting, centrifugation, and temperature variations required for the off-chip organoids, which may have further reduced the redox ratio values for the 0.3 µM group compared to their on-chip counterpart (Fig. 4c). Moreover, in contrast to the chip, the nutrient and reagent supply as well as the organoid growth rates are highly variable in a Matrigel dome and thus organoids’ metabolic activities and redox ratios could be variable depending on their location. Losing track of organoids when transferring them from their native culture to a secondary imaging platform is inevitable. Therefore, unlike in the chip, it is not possible to determine their native locations and set an additional control group to explain the lower redox ratios of Dox-treated off-chip organoids.

### On-chip immunofluorescence staining of organoids

Immunofluorescence staining is a powerful technique that allows investigation of complex biological processes in organoids through microscopic imaging^24^. For example, the cellular composition of intestinal organoids has been explored using specific antibodies, such as SOX9 and OLFM4 for intestinal stem cells, Lysozyme for Paneth cells, ChgA for enteroendocrine cells, MUC2 for goblet cells, and L-FABP for enterocytes^25,26^. Furthermore, many studies have used Ki67 antibodies to tag proliferative cells in both healthy intestinal organoids^4^ and colorectal cancer organoids^7,11^. These markers allow for the fluorescence visualization of various cell types and compartments, facilitating detailed studies of organoid biology. Recently, multiple studies have performed on-chip immunofluorescence^5–7,27^, DAPI (nuclei), and Phalloidin (F-actin) stainings^1^ on organoids grown using microfluidic devices, which illustrates the capabilities of such platforms for studying organoids.

Despite advancements in on-chip immunofluorescence staining, the limited diffusion of antibodies into Matrigel and organoids remains a challenge. Digestion of the Matrigel that encapsulates organoids is beneficial for promoting the diffusion, binding efficiency of the antibodies to the target antigens, and preventing non-specific binding with the Matrigel. On the other hand, dissolving the Matrigel causes other problems such as organoid drifting and blurry images in the absence of the Matrigel as a scaffold^8^. Therefore, organoids need to be suspended in mounting media and transferred to coverslips with imaging spacers for immobilization and imaging, a laborious procedure that is not conducive to high-throughput applications. On-chip staining eliminates the need for time-consuming centrifugation steps and reduces the risk of sample loss and morphology alterations caused by pipetting in conventional methods. Using the OrganoidChip+, we performed on-chip staining on 7-day-old organoids by first digesting the Matrigel, fixing with 4% PFA, and performing the necessary steps for staining (see Methods), all on the chip, followed by organoid immobilization in the TAs for high-resolution imaging.

Fixation with 4% PFA significantly reduces the adhesive properties of intestinal organoids. As a result, most organoids that had adhered to the glass during growth became detached and free after fixation, necessitating immobilization for high-resolution imaging. **Video S1** demonstrates the flow dynamics within the chip as the fixed and stained organoids move between the culture chamber and the TAs to distribute uniformly between all the TAs. Smaller organoids entered the immobilization chambers while the larger ones stayed at the staging chambers (**Figs. 5a-b**). Next, we tested the immobilization of organoids trapped in the TAs by subjecting them to flow pulses of 800 µL/min in both directions, as demonstrated in **Video S2**. The organoids remained secured in place within the TAs unaffected by the applied flow or movements of the chip, thus facilitating high-resolution imaging during fast stage transitions. While the shadow cast by the inlet glass reservoir decreased light exposure at areas close to the inlet during brightfield imaging, it did not influence the fluorescence imaging using markers such as DAPI or WGA (**Figs. 5c-d**). High-resolution images were captured from organoids in TAs 1 and 2 using a 0.75 NA, 20× objective following staining with DAPI and WGA (**Figs. 5e-f**). WGA staining highlighted the lumen of organoids by binding to the cell membrane^28^ and areas rich in mucus^29,30^. The strong WGA signal, observed within the lumen of these organoids indicates the presence of mucus-secreting cells, such as Goblet cells^31^. However, this hypothesis should be further investigated using Goblet cell-specific antibodies in future studies.

**Figure 5:**
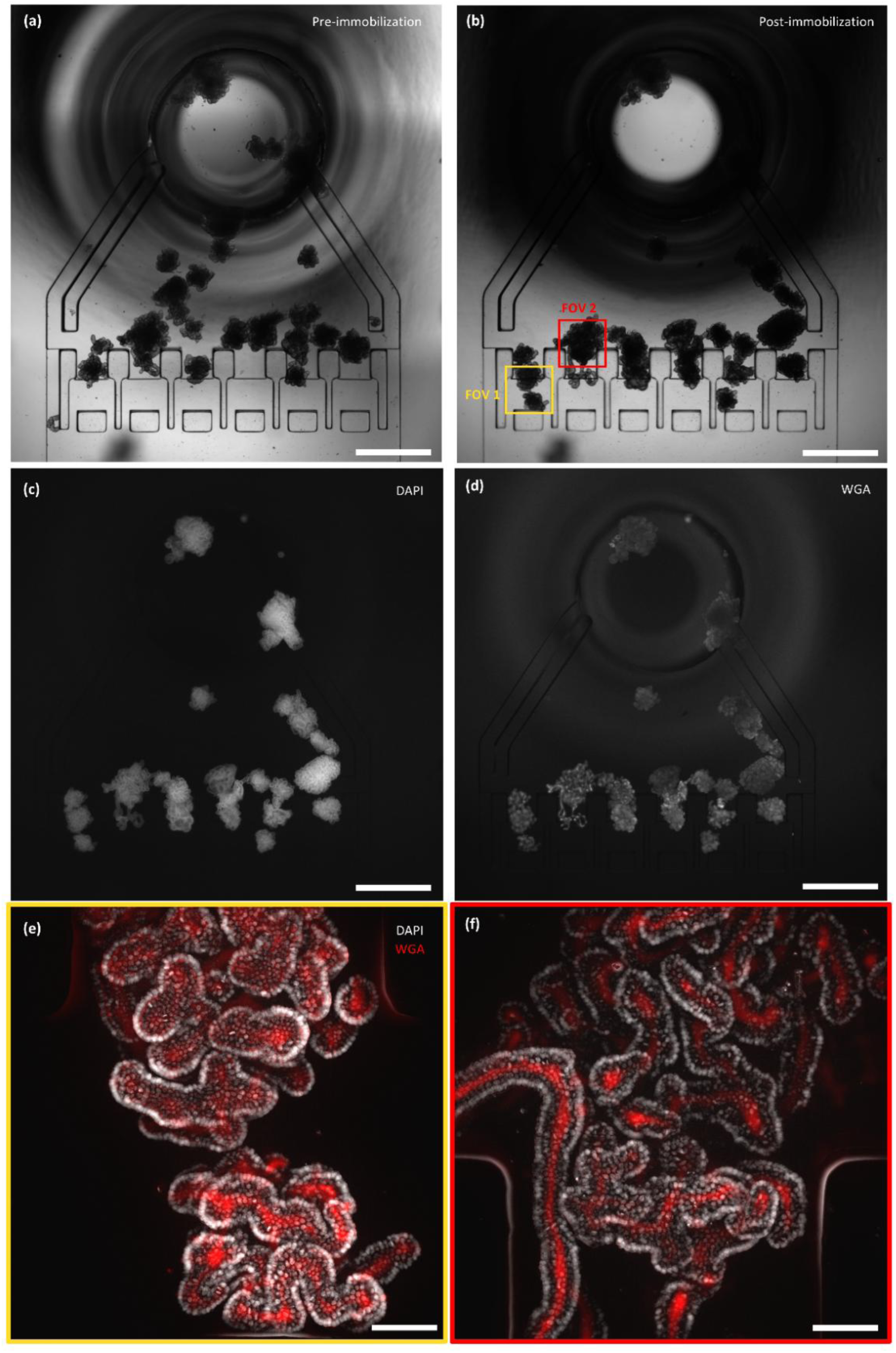
High-resolution images of organoids cultured, immobilized, and stained with DAPI and WGA, all performed within the same chip. (a-b) Brightfield images of the chip with organoids stained with WGA and DAPI on-chip, before and after immobilization, respectively. **(c-d)** Widefield fluorescence images captured with a 0.16 NA, 4× objective from DAPI and WGA, respectively. **(e-f)** High-resolution images captured with a 0.75 NA, 20× objective from organoids in FOVs 1 and 2 shown in (b). Images depict nuclei (DAPI, gray) and the intestinal lumen (WGA, red). Scale bars in (a-d) indicate 1 mm and those in (e-f) indicate 100 µm.

Figures 6 and **7** show immobilized organoids in the TAs, stained with ChgA and Ki67, respectively, and counterstained with DAPI and Phalloidin. Organoids were also counterstained with WGA (**Supplementary Fig. S6**). Both off- and on-chip organoids show similar expression levels of these key markers, which indicate healthy growth conditions in the chip (**Figs. 6c-i** & **Figs. 7b-i**). Furthermore, we counted the number of Ki67^+^ proliferative cells and ChgA^+^ enteroendocrine cells from confocal images, calculated their density, and compared them between organoids grown off- and on-chip (Figs. 6g and 7i). The densities of Ki67^+^ and ChgA^+^ cells were not statistically different between the off- and on-chip organoids (n = 6 organoids for each group). Enteroendocrine cells constitute a small fraction of the total cell population in the intestine, both *in vivo* and in organoid models^25,27^. The presence of the rare ChgA^+^ enteroendocrine cells within the chip highlights its ability to replicate relevant *in vitro* microenvironment (white arrows in Figs. 6e-f and 6i). Moreover, the Ki67^+^ cells are observed in many areas, including regions near the tip of the crypts (white arrows in Figs. 7c-d and 7h), which indicate pertinent cellular composition regarding proliferative stem and progenitor cells in our organoids^4^.

**Figure 6:**
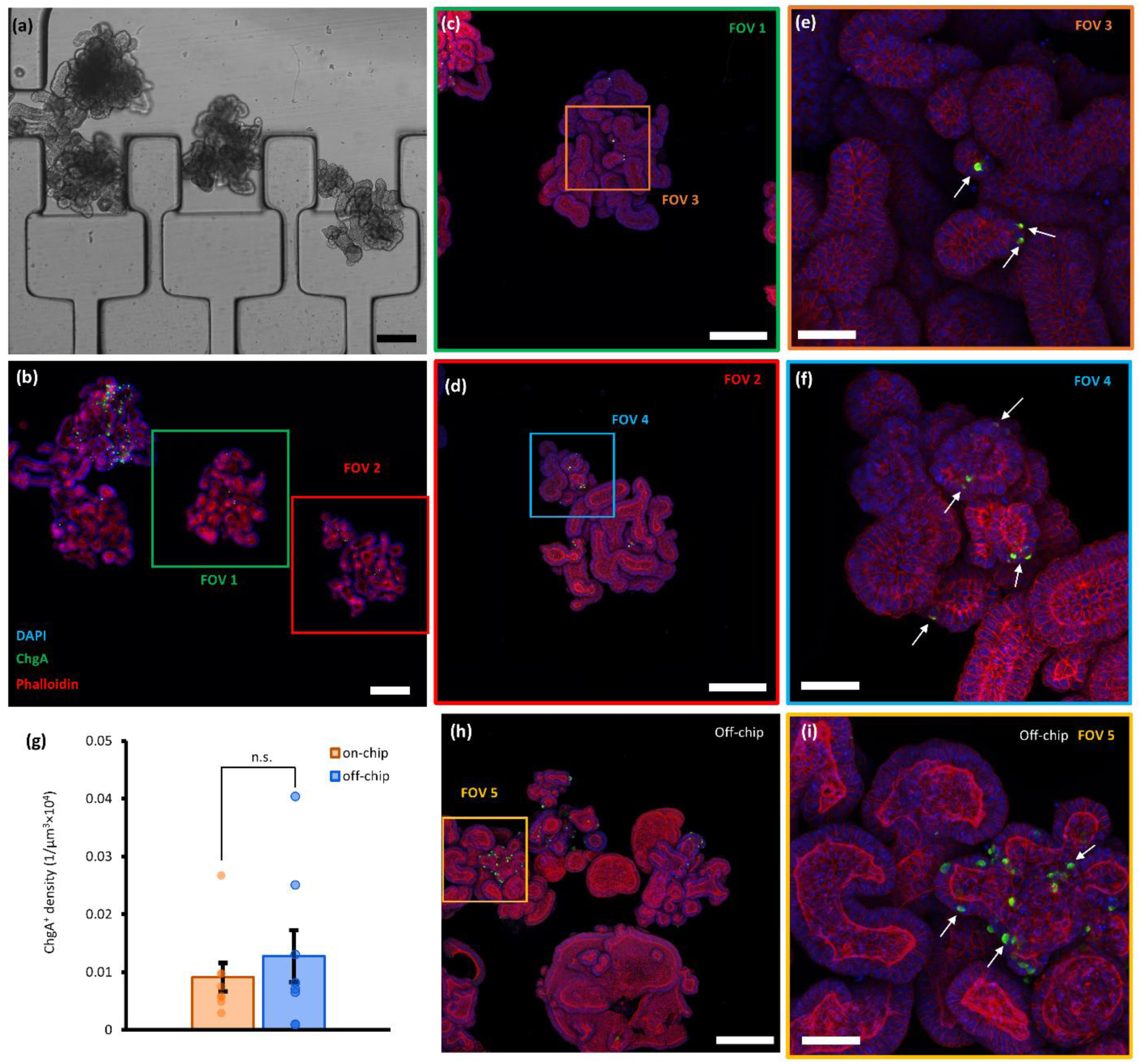
Fluorescence confocal images of organoids captured following growth and immunofluorescence (ChgA – green), DAPI (blue), and phalloidin (red) staining off- and on-chip. (a-b) Brightfield and widefield fluorescence images of organoids immobilized in the SCs or ICs. **(c-d)** FOVs 1 and 2 were recaptured using confocal microscopy (0.3 NA, 10×). **(e-f)** Confocal images (0.6 NA, 40×) of FOVs 3 and 4 shown in (c) and (d). **(g)** Compares the density of ChgA^+^ cells between off- and on-chip organoids with no statistically significant differences. n = 7 FOVs per condition, captured with a 0.6 NA, 40× objective (two-way *t*-test, *p*- value = 0.22). **(h)** Image from off-chip organoids (0.3 NA, 10×). **(i)** FOV 5 was recaptured using a 0.6 NA, 40× objective. White arrows point out the ChgA^+^ enteroendocrine cells. Scale bars represent 200 µm in (a-d) and (h), and 50 µm in (e-f) and (i).

**Figure 7:**
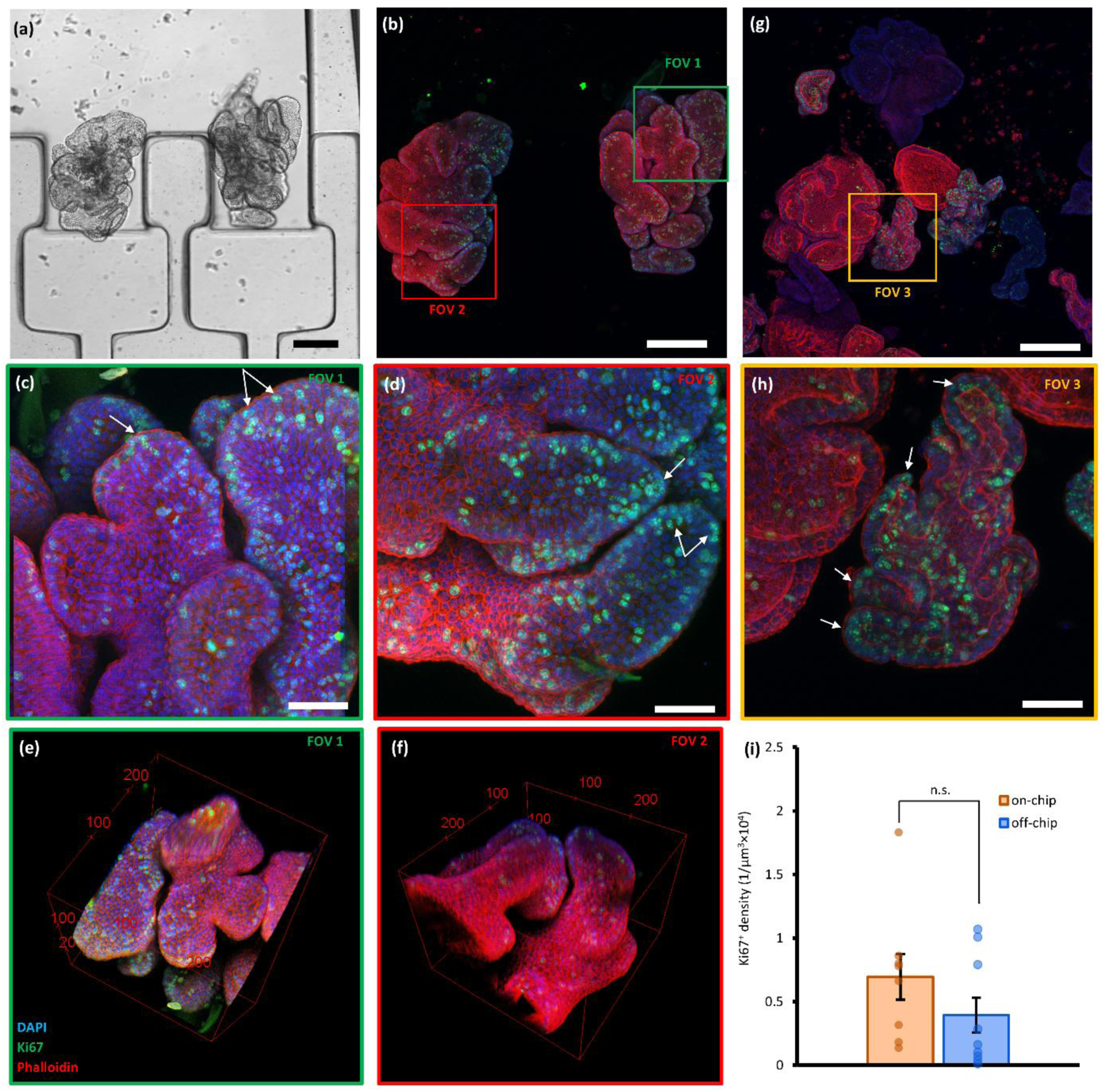
Fluorescence confocal images of organoids captured following growth and immunofluorescence (Ki67 – green), DAPI (blue), and phalloidin (red) staining off- and on-chip. **(a)** Brightfield (0.16 NA, 4×) and **(b)** confocal fluorescence images (0.3 NA, 10×) of organoids, stained and immobilized in TAs in the chip following their 7-day-long culturing in the chip. **(c)** FOV 1 and **(d)** FOV 2, were recaptured using a 0.6 NA, 40× objective. **(e-f)** 3D reconstruction of FOVs 1 and 2 using z-images captured with a step size of 3 µm. The bounding boxes serve as scale bars with dimensions in microns. **(g)** Image from off-chip organoids (0.3 NA, 10×). **(h)** FOV 3 was recaptured using a 0.6 NA, 40× objective. White arrows point out the Ki67^+^ proliferative cells located at the crypts. **(i)** Plot of Ki67^+^ cells densities observed both off- and on-chip organoids, showing no statistically significant differences. n= 8 FOVs per condition, captured with a 0.6 NA, 40× objective (two-way *t*-test, *p*-value = 0.52). Scale bars represent 200 µm in (a), (b), and (g), and 50 µm in (c), (d), and (h).

High-resolution confocal images from two field-of-views (FOVs) in **Figs. 7c-d** (**Videos S6-S7**) were used to reconstruct the 3D view of organoids shown in **Figs. 7e-f** (**Videos S8-S9**), respectively. Imaging was performed up to a depth of 250 µm within the organoids, with deeper regions exhibiting increased light scattering, which reduced the signal-to-background ratio (**Videos S6-S7**). Nevertheless, the 3D reconstruction effectively revealed lumen morphology and provided a better understanding of the spatial distribution of target cells.

## Conclusion

In this study, we developed an organoid culturing, staining, immobilization, and imaging all-in-one device that enables transferless, blur-free, fast, and HCI of immobilized organoids. We added perfusion channels to our previously designed OrganoidChip^8^ to facilitate on-chip culturing of ASOs over 7 days with growth rates equal or higher than conventional Matrigel dome cultures. We implemented viability and redox ratio assays, and probed ChgA and Ki67 markers with immunofluorescence while organoids were counterstained with WGA, Phalloidin, and DAPI. As proof-of-concept, HCI was performed using high-resolution imaging modalities suited for each assay, all inside the chip without transferring organoids to any secondary platform. This end-to-end and transferless organoid culture, staining, and imaging process hinders sample loss, protects from pipetting shear, prevents losing track of organoids, and minimizes the number of preparation steps used in existing methods.

To verify the health of organoids grown on-chip, we conducted viability staining using Calcein AM and EthD-1 and found that their viability closely matched that of off-chip organoids. Next, we treated 5-day-old organoids with Dox for 48 hours, both off- and on-chip, followed by label-free, two-color, two-photon redox ratio imaging. The results revealed that off-chip organoids exhibited lower redox ratio levels at reduced Dox concentrations relative to their on-chip counterpart. This difference might be attributed to the shear stress and temperature variations encountered during pipetting, centrifugation, and additional Matrigel embedding steps in off-chip handling of organoids. Also, the highly variable nutrient and reagent diffusion in dome cultures may have resulted in lower redox ratios, particularly in organoids located near the center of the Matrigel dome. Our platform overcomes these limitations by maintaining organoids on-chip throughout the entire culturing, staining, and imaging process, ensuring traceability and minimizing handling-induced variability.

End-to-end tracking of organoid development from growth through compound treatment all the way to immunostaining is crucial for identifying key parameters involved in biological processes such as stem cell differentiation, proliferation, and changes in cellular composition. Such studies need platforms that support transferless organoid imaging, which is vital for performing comprehensive morphological and molecular biomarker analyses. To this end, we conducted immunofluorescence, Phalloidin, WGA, and DAPI staining of organoids followed by immobilization and high-resolution confocal imaging, all within the chip. Because organoids fixed with 4% PFA lose their adherent properties, they can freely move during fast imaging sessions and upon any disturbance to the platform, without adhering to the glass or PDMS surface. As opposed to the currently existing platforms for ASO culturing and imaging, the chip has the advantage of immobilizing these organoids at predetermined locations, within the TAs, to facilitate high-resolution and blur-free imaging in a fully automated manner. Our results demonstrated close agreement between off- and on-chip expression levels of ChgA and Ki67 markers, indicating similar volumetric densities of enteroendocrine and proliferative cells, respectively. We successfully performed 3D reconstruction of stained organoids, which can be valuable for understanding the spatial distribution of specific cell types and the mechanisms of action in bacterial infectious diseases, in future studies^32^.

These advancements were made possible by the specific design of the OrganoidChip+ that fully meets the requirements for successful HCI of ASOs. The chip’s design encompasses several key features that offer distinct advantages over existing technologies. Most notably, it facilitates full immobilization of organoids after their release from Matrigel by anchoring them within six trapping areas to prevent any movement and blur during imaging. The trapping areas also provide pre-determined locations for organoids, allowing for automatic positioning of the microscope stage for rapid image acquisition. Another critical design consideration for successful HCI was ensuring optical and geometrical compatibility with high-resolution microscope objectives. To this end, the chip incorporates a thin glass substrate and a short-height culture chamber to minimize the required number of z-stack images. Furthermore, we chose channel dimensions that are compatible with cost-effective, large-scale fabrication methods such as injection molding of plastic materials. The use of plastic instead of PDMS will eliminate the absorptive properties of PDMS that can interfere with test results, especially when working with small hydrophobic molecules^33^.

While the OrganoidChip+ has demonstrated strong performance, a few limitations are being addressed in upcoming studies. First, the current chip design supports only organoids within a specific size range (250 – 550 µm), which can be reliably immobilized within the trapping areas. However, the geometry of the chip, especially those of the TAs, can readily be modified to accommodate a broader range of organoid sizes or types. Second, the current data using canine-derived organoids serves as a proof of concept supporting the use of this technology for preclinical toxicology assessment of therapeutic drug candidates, with the goal of reducing reliance on live animal testing in alignment with the 3Rs (Replacement, Reduction, and Refinement). This platform may also support regulatory submissions in veterinary drug development, particularly when the dog is the intended target species. Moving forward, scaling this technology to human-derived organoid models will further enhance translational relevance by enabling more accurate prediction of human responses to candidate therapeutics.

In future developments, OrganoidChip+ can be scaled up into a 24- or 48-well format and fabricated in standard SBS dimensions to support high-throughput studies. Interconnecting multiple chips with different ASO types could enable the creation of a multi-organ-on-chip system with automated fluid handling and imaging capabilities. Moreover, the current design allows for easy retrieval of the conditioned media from the outlet reservoir for myriads of downstream analyses such as protein profiling^34^ and metabolic analysis^35^ that are crucial for understanding the cross-talk between different ASOs. With these enhancements fully implemented, we anticipate that the OrganoidChip+ will emerge as a highly competitive platform for organoid culturing, staining, immobilization, and HCI. Its ability to support end-to-end analysis of organoids within a single, integrated system uniquely positions it as a valuable tool for both basic research and translational applications.

## Methods and materials

### Device fabrication

We fabricated the microfluidic chip as previously described^8^. Briefly, we first fabricated a three-layer mold using photolithography with negative photoresist SU8-2100, followed by soft lithography with polydimethylsiloxane (PDMS). The three photoresist layers, with heights of 100, 190, and 260 µm, were coated on a silicon wafer substrate to produce channel heights of 100, 290, and 550 µm, respectively. First, a 6-inch silicon wafer (STK9671-1, Nova Electronic Materials) was dehydrated on a hot plate at 150 °C for 20 minutes, then placed onto a spin coater. To achieve the desired layer thickness, 6 mL of the photoresist was dispensed at the center of the wafer and spun. After spinning, the wafer was removed and soft-baked, then allowed to cool to room temperature. It was subsequently exposed to UV light through a high-resolution mylar mask tailored to the specific layer. The photoresist was developed after the third UV exposure (for more details on the process times and temperatures, see **Supplementary Table S1**). To facilitate the easy removal of the mold from the PDMS, the mold was treated with a silanization process using (TRIDECAFLUORO-1,1,2,2-TETRAHYDROOCTYL) TRICHLOROSILANE (Gelest, Inc.) for 72 hours.

For the soft lithography process, the base polymer and curing agent were mixed in a 10:1 (w/w) ratio, degassed in a vacuum chamber, and then poured onto the mold. It was then oven-baked at 80 °C for 6 hours. After curing, the PDMS was peeled off, punched using a gauge 11 punch (CR1200945N11R1, Syneo, LLC) for inlet and outlet holes, and bonded to a #1.5 cover glass using oxygen plasma treatment to form the final chip.

### Ethical Animal Use

This colon sample was obtained from a necropsy case for an unrelated research project. The collection of samples from dogs was conducted in accordance with Standard Operating Procedures (SOPs) approved by the Marshall BioResources’ Institutional Animal Care and Use Committee. All methods were performed in accordance with the USDA Animal Welfare Regulations.

### Organoid culture and maintenance

Intestinal crypts were obtained from an adult healthy canine intestine as previously described^36^. In this study, the intestinal organoid culture and maintenance were performed according to our previous publication with small modifications^8^. Briefly, the organoid culture was established by thawing a frozen vial containing organoids in a 37 °C water bath. The content was added to 4 mL of DMEM (12634010, Advanced DMEM/F-12, Gibco^TM^) and centrifuged at 100 g for 5 minutes at 4°C. The supernatant was removed, 90 µL of ice-cold Matrigel was added to the pellet on ice with a 200 µL pipette tip and pipetted up and down to homogenously mix the organoid-Matrigel suspension. 25 µL of the suspension was dispensed per well in a 24-well plate to form the Matrigel domes. The plate was incubated at 37 °C for 10 minutes for the gel to solidify. Then, 0.5 mL of the Complete medium with Growth Factor (CMGF^+^)^8^ was added to each well containing the organoids and incubated (See **Supplementary table S2** for the CMGF^+^ preparation protocol). The CMGF^+^ was refreshed every 2 days until clean-up or passage.

Organoids were ready for a clean-up between Days 3 to 4 and for a passage between Days 7 to 10, depending on their size and density in the Matrigel dome. The clean-up and passaging protocols were previously described^8^. Briefly, the used CMGF^+^ in each well was replaced 0.5 mL of 4°C Corning® Cell Recovery Solution (CRS) and pipetted multiple times to break the Matrigel dome. The resulting suspension was centrifuged at 100 g for 5 minutes at 4°C and then the supernatant was removed. For a clean-up, desired amount of ice-cold Matrigel was added to the pellet, pipetted and seeded in a 24-well plate as 25 µL domes in each well. For passaging, 1 mL of TrypLE Express (Cat# 12-604, ThermoFisher Scientific) was dispensed to the pellet, pipetted, and incubated for 6 minutes at 37 °C and 5% CO_2_. Then, we pipetted the mixture a few times and added 4 mL of DMEM to stop the dissociation. We centrifuged the suspension, removed the supernatant and added 1 mL of DMEM. The suspension was pipetted multiple times for homogenization and immediately used to count the number of clumps using a standard Hemocytometer slide. After counting the number of clumps, the necessary amount of the suspension was withdrawn and centrifuged at 100 g for 5 minutes at 4°C. To yield the final seeding density of 1.5 × 10^5^ clumps/mL of Matrigel, the necessary amount of 4°C Matrigel was added to the pellet and seeded in the wells as 25 µL domes.

### Organoid culture on-chip

After plasma bonding the PDMS with the cover glass, the chips were kept under a sterile laminar hood wrapped in aluminum foil for a week before use. On the day of seeding, the chips were exposed to UV inside the laminar hood for 30 minutes. Next, the cell-Matrigel suspension was prepared at the same density as in dome culture and injected into the culture chamber of the chip using a bent Luer stub, as shown in **Figs. 1b-c** (Blunt needle, gauge 25, Instech Laboratories). The Luer stub was bent at ∼ 2 mm distance from its tip to achieve a 90-degree angle. The Luer stub was connected at the tip of a p50 pipette (VWR, Inc.) and sealed with Teflon tape. For seeding, we withdrew 5 µL of the suspension via the bent needle, inserted the tip of the needle into the culture chamber and dispensed slowly until the entire culture chamber was filled (**Figs. 1b-c**). The chip was placed in a petridish and transferred to the incubator for 10 minutes to solidify the Matrigel. Next, we mounted sterile glass reservoirs (Precigenome LLC) at the inlet and outlet of each chip and added 200 µL of CMGF^+^ to the inlet reservoir (**Supplementary Fig. S1a**). The openings of the reservoirs were sealed with a thin parafilm to avoid the introduction of debris and contamination as well as preventing any moisture loss during imaging sessions. The culture medium was refreshed every 48 hours by removing the used medium from the inlet and outlet reservoirs and adding 200 µL of the fresh CMGF^+^ to inlet reservoir.

### On-chip Matrigel digestion

First, the culture medium in the reservoirs were removed and 200 µL of cold CRS was added to the inlet reservoir. After 10 minutes of incubation at 4°C, the CRS from the reservoirs were removed and another 200 µL of cold fresh CRS was added to the inlet reservoir. The chip with the CRS flowing from the inlet to the outlet reservoir was kept at 4°C for an additional 10 minutes for the full digestion of Matrigel. Next, we removed the CRS from the reservoirs and added 200 µL of the desired reagent to the reservoirs for the subsequent process.

### On-chip organoid immobilization

The experimental protocol for the immobilization process and the fluidic setup have been detailed in our prior publication^8^. Briefly, the fluidic setup consisted of an air pressure controller (ITV0010-3UBL, SMC) that is connected to two large reservoirs containing a working fluid via two solenoid valves. We either used DMEM for live/dead viability staining or PBS for chips with immunostained organoids as the working fluid. The architecture of the tubing and the two solenoid valves formed a fluidic H-bridge to facilitate the flow of liquid in both directions inside the chip. A DAQ card (NI USB-6009) was controlled via a LabVIEW program to regulate the flow rate and the solenoid valve functions.

For organoid immobilization after Matrigel digestion, 200 µL of the working fluid was added to each of the chip reservoirs. Then, the chip reservoirs were carefully connected to the inlet and outlet tubings of the fluidic setup without introducing any bubbles. Organoids were guided into the TAs using flow rates of up to approximately 160 µL/min, corresponding to an upstream pressure of ∼ 0.1 bar. In cases where higher drag forces were required to overcome adverse friction and facilitate organoid entry to the TAs, brief pulses of higher flow rates up to 800 µL/min (0.5 bar) for less than 1 second were applied. Using this immobilization scheme, minimal to no cell shedding was observed from the live organoids, suggesting that the applied shear stress did not cause any physical damage to them.

### Live/dead staining and analysis

We stained the immobilized organoids on-chip using Calcein AM (Invitrogen™), Ethidium Homodimer-1 (EthD-1) (Invitrogen™), and Hoechst (33342, ThermoFisher Scientific Inc.) to label live cells, dead cells, and the cell nuclei, with concentrations of 2, 4, and 2 µM, respectively. All three dyes were prepared in CMGF^+^ immediately before use and pooled together for simultaneous administration. For on-chip staining, organoids were first immobilized as mentioned in the previous section, then, DMEM was removed from the reservoirs and replaced with 200 µL of the dye solution. After 30 minutes of incubation at 37°C and 5% CO_2_, the used dye solution in the reservoirs were removed and fresh dye solution was added to the inlet reservoir. Next, the reservoir openings were sealed with thin parafilm, and the chip was incubated at 37°C and 5% CO2 for another 30 minutes prior to imaging.

For off-chip viability staining, multiple 1 mL pipette tips were cut at the tip and rinsed in cell culture grade water twice. Then, the culture medium in each well was replaced with 0.5 mL of cold CRS. The cut tips were used to pipette the CRS up and down to break the Matrigel domes and transfer the content to a 15 mL falcon tube without imposing excessive shear stress on organoids. The falcon tube was centrifuged at 100 g for 1 minute, the supernatant was removed, and the pellet was suspended in the dye solution. The content was added to an 8-well µ-slide with #1.5 glass bottom (80827, Ibidi Inc.) and incubated at 37°C and 5% CO_2_ for 1 hour prior to imaging.

For viability analysis, we created a mask for each organoid using the minimum intensity projection image and thresholding to obtain the boundaries of organoids^8^. Then, we used the maximum projection image of the green and red channels and calculated the mean fluorescence intensity inside the masks corresponding to each organoid (live: S_GO_ and dead: S_RO_). To calculate the viability (V), we primarily adhered to the manufacturer’s protocol and referenced previous studies^8,37^. Additionally, we incorporated a normalization step to enhance the accuracy of the viability calculations. Specifically, the ratio of live cells to the total number of cells (live and dead) was calculated for each organoid by normalizing the mean fluorescence intensity for each channel using the mean intensity of a single live cell (S_GC_) and a single dead cell (S_RC_). The viability was then determined using the following formula:

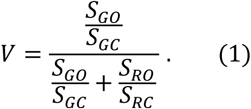

This normalization step ensured consistency across samples and minimized variability caused by differences in imaging conditions or staining efficiency.

### Automation and widefield imaging

Widefield and fluorescence imaging were conducted using an ORCA-Flash4.0 V2 Digital CMOS camera (Hamamatsu Photonics) in combination with an IX73 Inverted Microscope (Olympus) equipped with GFP, TexasRed, and DAPI filter cubes. Automated imaging and positioning of well plates and the chip on the IX73 microscope were achieved using a PZU-2000 automated stage and an ASI MS-2000-WK stage controller. Z-imaging was performed using a 0.16 NA, 4×, a 0.3 NA, 10×, and a high-resolution 0.75 NA, 20× objective (Olympus) with z-step sizes of 20, 10, and 3 µm, respectively. The imaging area covered all the TAs and the entire culture chamber. The imaging setup was automated via a custom-built LabVIEW (NI, USA) program. All image processing was carried out in Fiji (NIH).

### Label-free, two-color, two-photon redox ratio imaging

For off-chip redox ratio imaging, we first retrieved the organoids from the Matrigel domes by adding 0.5 mL of cold CRS per well and 1 mL pipette tips that were cut at 1 mm distance from their tip. The domes were triturated a few times without imposing high shear stress on organoids. The organoids were centrifuged in a 15 mL tube at 100 g for 3 minutes. The pellet was suspended in ice-cold Matrigel and dispensed as 40 µL droplets at the center of a cover glass. Previously, two pieces of tapes, each a stack of multiple lab tapes with a total thickness of 550 µm, were adhered to either side of the place where droplet was dispensed. Another cover glass was mounted on top of the droplet to create an organoid sandwich with a gap height of 550 µm. The organoid sandwich was incubated at 37°C and 5% CO_2_ for 10 minutes prior to imaging.

For on-chip redox ratio imaging, we did not need to digest the Matrigel. Simply, the reservoirs were detached from the chips, and a cover glass was positioned over the inlets and outlets to create a sealed environment. Next, the chips and the organoid sandwiches were transferred to the two-photon microscope for imaging.

We used a Ti:sapphire laser (Mai Tai, Spectra-Physics) with 100 fs pulse duration and 80 MHz repetition rate to excite the samples. The laser beam was coupled to an in-house-built two-photon microscope, optimized for increased signal collection efficiency^19^. The beam was expanded by a factor of 4.1 to fill the back aperture of a 0.75 NA 20x (Nikon Olan Apo) objective. On the emission path, we used a short-pass filter (BrightLine® 720/SP, Semrock) to block off the excitation beam and an additional fluorescence filter (BG39, Schott) to collect NADH and FAD signals. Collected fluorescence was sensed by a PMT (H10770PA-40), which interfaced with a low-noise preamplifier (SR570). We adjusted the wavelength of the laser source at 745 nm and 860 nm to excite NADH/FAD and FAD molecules, respectively, and thereby to perform redox ratio analysis. First, the organoid surface was identified at the excitation wavelength of 745 nm and then, organoids were imaged every 20 µm, starting at a depth of 10 µm up to a depth where signal-to-background ratio was sufficiently high to resolve features. At each depth, 20 images were collected at a frame rate of 1 Hz. After reaching the maximum depth, the excitation wavelength was switched to 860 nm and the images were captured in reverse, from the deepest regions back to the organoid surface using the same frame settings. See **Supplementary Fig. S5** and **Eqs. S6-S12** for detailed two-photon image analysis and redox ratio calculations.

### Immunofluorescence staining

The preparation of solutions began with the preparation of a PBS solution containing 0.1% (v/v) Tween- 20 (PBT) by adding 20 µL of Tween-20 to 2 mL of PBS (10×) (Cat# 14200, ThermoFisher Scientific) and 18 mL cell culture grade water (Cat# W3500, Sigma-Aldrich). Organoid washing buffer (OWB) was prepared according to Martinez-Ordoñez *et al.* by combining 0.5 mL of 10% Triton X-100 in PBS (1×), 50 µL of 10% (w/v) SDS in water, and 0.1 g of bovine serum albumin (BSA) with 49.5 mL of PBS (1×), resulting in final concentrations of 0.1% Triton X-100, 0.01% SDS, and 0.2% BSA^38^. A glycine solution was prepared by dissolving glycine in PBS (1×) to a final concentration of 300 mM. Also, several 1 mL pipette tips were modified by cutting the last 1 mm and pre-rinsing with Anti-Adherence Rinsing Solution (STEMCELL Technologies) to avoid excessive shear stress and adherence of organoids to the pipette tips. The primary antibodies used consist of Ki67 (FITC, 50-112-9478, Fisher scientific) and ChgA (20085, ImmunoStar Inc.). For ChgA, we used the Alexa Fluor™ Plus 488 secondary antibody (A32731, ThermoFisher Scientific). The cells were counter stained with DAPI (62248, ThermoFisher Scientific) and Rhodamine Phalloidin (R415, ThermoFisher Scientific) or WGA (29023, Biotium) both off- and on-chip.

For fixing organoids off-chip, the organoids were first recovered from Matrigel in the well plate by replacing the culture medium with 0.5 mL of cold CRS. The domes were triturated gently several times, and the contents of the wells were transferred to a 15 mL conical tube. The tube was then filled with 10 mL of ice-cold PBS (1×) and centrifuged at 4°C, 100 g for 3 minutes. The supernatant, containing most of the Matrigel, was aspirated, and the organoid pellet was fixed by resuspending it in 1 mL of 4% PFA (Santa Cruz Biotechnology), followed by incubation at 4°C for 1 hour. In the middle of the fixation period, the organoids were gently resuspended once.

For on-chip organoid fixation, Matrigel was first digested, and organoids were immobilized as described in previous sections. Then, the PBS in the reservoirs was removed, 200 µL of 4% PFA was added to the inlet reservoir and the chip was incubated at 4°C for a total of 1 hour. The 4% PFA solution was refreshed in the middle of the incubation time to promote the fixation process.

After fixation, the 4% PFA was removed and ice-cold PBT was added to the tube (10 mL) and the chip (200 µL) and incubated at 4°C for 10 minutes. Next, the PBT was removed and organoids in the tube and the chips were washed twice for 20 minutes each with the glycine solution. For samples that were counter-stained with WGA, we prepared a 5 µg/mL WGA in PBS (1×) solution and carried out an overnight treatment at 4°C. To remove the liquids from the tube, we centrifuged the tube at 100 g for 3 minutes followed by aspirating the supernatant and for the chips we simply removed the liquid from the reservoirs.

For blocking and permeabilization, the glycine solution was removed from both the tube and the chip. Organoids were treated with OWB both off- and on-chip at 4°C for 15 minutes. In parallel, the primary antibody solution was made in OWB (ChgA 1:100, Ki67 1:100). Then, the OWB was replaced with the primary antibody solution. The organoids in the tube were transferred to a 96-well plate and placed on an orbital shaker at 80 rpm (Orbi-Shaker™ JR., Benchmark Scientific Inc.). The chips were placed on a Nutating mixer (Cat# 05-450-213, Fisher Scientific). Both off- and on-chip experimental samples were incubated at 4°C for 24 hours.

Next, both off- and on-chip organoids were washed with OWB for 3 × 2 hours at 4°C. The secondary antibody solution (2 µg/mL) was prepared in OWB in combination with DAPI (0.67 µg/mL) and Phalloidin (1:40). The OWB in the wells and the chips was replaced with the secondary solution. The plate and the chips were covered with aluminum foil and incubated at 4°C for 24 hours with shaking and tilting. Finally, organoids were washed with OWB for 3 × 2 hours at 4°C. Organoids in the 96-well plate were transferred to an 8-well µ-slide (Ibidi Inc.) for imaging. On-chip organoids were imaged as being immobilized on the chip.

### Confocal imaging

Confocal imaging of the immunostained organoids was performed using a Leica SP8 microscope. Z-stacks of Ki67 (excitation: 488 nm, bandpass: 492–541 nm), ChgA (excitation: 488 nm, bandpass: 505–539 nm), Phalloidin (excitation: 552 nm, bandpass: 560–780 nm), WGA (excitation: 552 nm, bandpass: 550–780 nm) and DAPI (excitation: 402 nm, bandpass: 410–489 nm) were acquired using 10×, 0.3 NA and 40×, 0.6 NA objectives with step sizes of 5 µm and 3 µm, respectively. All image processing was carried out in Fiji.

### Statistical analysis

We used Microsoft Excel (Microsoft 365 MSO, Version 2410) to perform data and statistical analysis. A significance level of 0.05 was used for hypothesis testing. After checking for equality of variances using an *F*-test, one-tail or two-tail *t*-tests were used for group comparisons with “*”, “**”, “***”, and “n.s.” indicating *p* < 0.05, *p* < 0.005, *p* < 0.0005, and *p* > 0.05, respectively. The error bars represent the standard error of the mean (SEM) unless otherwise specified.

## Conflict of interest

A.B., K.M., S.M., E.H., and A.H. are listed as inventors on a pending U.S. patent application (PCT/US24/13330) directly related to the platform described in this work, which has been licensed to vivoVerse, LLC. Additionally, A.B., S.M., and E.H. hold equity in vivoVerse, LLC. These disclosures have been reviewed and managed according to institutional policies addressing potential conflicts of interest in research. K.A. is a co-founder of LifEngine Animal Health and 3D Health Solutions and serves as a consultant for Ceva Animal Health, Bioiberica, LifeDiagnostics, Antech Diagnostics, Deerland Probiotics, Christian Hansen Probiotics, Purina, and Mars. J.P.M. is a co-founder of LifEngine Animal Health (LEAH) and 3D Health Solutions and serves as a consultant for Ceva Animal Health, Ethos Animal Health, LifEngine Animal Health and Boehringer Ingelheim. C.Z. is the Director of Research and Product Development at 3D Health Solutions.

## Data availability

Data for this article, including figures, codes, and videos are available at box.com at https://utexas.box.com/s/88k85dypn8s5ozzt3uq2xrxibmjxirq.

## Acknowledgments

We thank Dr. Marissa N. Rylander for providing access to their confocal microscope and Daniel Wu for his valuable input during the development of automation. This work was partially supported by the National Institutes of Health (NIH) grant # R43-ES029890 from the National Institute of Environmental Health Sciences (NIEHS) that was awarded to vivoVerse, LLC.

## CRediT authorship contribution statement

**Khashayar Moshksayan**: Conceptualization, Methodology, Investigation, Software, Formal analysis, Data curation, Visualization, Validation, Resources, Supervision, Writing – original draft, Writing – review & editing. **Rishika Khanna**: Investigation, Resources, Formal analysis, Data curation. **Berk Camli**: Investigation, Methodology, Software, Resources. **Zeynep Nevra Yakay**: Methodology, Software, Formal analysis, Data curation. **Anirudha Harihara**: Methodology, Resources, Writing – review & editing. **Christopher Zdyrski**: Methodology, Resources, Writing – review & editing. **Sudip Mondal**: Conceptualization, Funding acquisition, Writing – review & editing. **Evan Hegarty**: Conceptualization, Methodology. **Jonathan P. Mochel**: Methodology, Resources, Writing – review & editing. **Karin Allenspach**: Methodology, Resources, Writing – review & editing. **Adela Ben-Yakar**: Conceptualization, Investigation, Resources, Funding acquisition, Supervision, Project administration, Writing – original draft, Writing – review & editing.

## Supplementary Information

**Video S1**

Immobilization of fixed organoids within the TAs using a controlled flow. Organoids are redistributed in the culture chamber multiple times by reversing the flow direction to trap organoids in all the TAs.

**Video S2**

Shows the chip with the immobilized organoids in **Video S1** being exposed to large flow rates (up to 800 µl/min) in both directions within the chip. As the flow rate and direction are changed during the disturbation, organoids reside immobilized within the TAs, showing the chip’s immobilization capabilities. The flow direction is evident by the direction of the particle or cell movements in the chip.

**Videos S3-S5**

Label-free, two-color, two-photon imaging of organoids at increasing depths. The images on the left and right were captured using 745 nm and 860 nm wavelengths, respectively. **Videos S3-S5** correspond to the on-chip vehicle control, 0.3 µM, and 2 µM Dox-treated organoids, respectively. The dimension of the FOV is approximately 400 µm.

**Videos S6-S7**

High-resolution confocal z-imaging using a 0.6 NA, 40× objective from organoids shown in **Figs. 7c-d**. The dimensions of the FOV are 291 µm by 291 µm.

**Videos S8-S9**

3D reconstruction of organoids created from the confocal images shown in **Videos S6-S7**. A snapshot of **videos S8-S9** are shown in **Figs. 7e-f**, respectively.

## Notes

https://utexas.box.com/s/88k85dypn8s5ozzt3uq2xrxibmjxirqm

## References

1 Schuster, B. et al. Automated microfluidic platform for dynamic and combinatorial drug screening of tumor organoids. Nature Communications 11, 5271, doi:10.1038/s41467-020-19058-4 (2020).

2 Hu, Y. et al. Lung cancer organoids analyzed on microwell arrays predict drug responses of patients within a week. Nature Communications 12, 2581, doi:10.1038/s41467-021-22676-1 (2021).

3 Lohasz, C., Frey, O., Bonanini, F., Renggli, K. & Hierlemann, A. Tubing-Free Microfluidic Microtissue Culture System Featuring Gradual, in vivo-Like Substance Exposure Profiles. Frontiers in Bioengineering and Biotechnology 7, doi:10.3389/fbioe.2019.00072 (2019).

4 Park, S. E. et al. Geometric engineering of organoid culture for enhanced organogenesis in a dish. Nature Methods 19, 1449–1460, doi:10.1038/s41592-022-01643-8 (2022).

5 Geyer, M. et al. A microfluidic-based PDAC organoid system reveals the impact of hypoxia in response to treatment. Cell Death Discovery 9, 20, doi:10.1038/s41420-023-01334-z (2023).

6 Miller, C. P. et al. Therapeutic targeting of tumor spheroids in a 3D microphysiological renal cell carcinoma-on-a-chip system. Neoplasia 46, 100948, 10.1016/j.neo.2023.100948 (2023).

7 Beghin, A. et al. Automated high-speed 3D imaging of organoid cultures with multi-scale phenotypic quantification. Nature Methods 19, 881–892, doi:10.1038/s41592-022-01508-0 (2022).

8 Moshksayan, K. et al. OrganoidChip facilitates hydrogel-free immobilization for fast and blur-free imaging of organoids. Scientific Reports 13, 11268, doi:10.1038/s41598-023-38212-8 (2023).

9 Moshksayan, K. et al. A microfluidic chip with immobilization chambers for cardiac organoid imaging. Vol. 12374 PWB (SPIE, 2023).

10 Kim, S.-O., Kim, J., Okajima, T. & Cho, N.-J. Mechanical properties of paraformaldehyde-treated individual cells investigated by atomic force microscopy and scanning ion conductance microscopy. Nano Convergence 4, 5, doi:10.1186/s40580-017-0099-9 (2017).

11 Wang, Z. et al. Rapid tissue prototyping with micro-organospheres. Stem Cell Reports 17, 1959–1975, 10.1016/j.stemcr.2022.07.016 (2022).

12 Promega, C. CellTiter-Glo® Luminescent Cell Viability Assay. Technical Bulletin (2009).

13 Alam, S. R. et al. Investigation of Mitochondrial Metabolic Response to Doxorubicin in Prostate Cancer Cells: An NADH, FAD and Tryptophan FLIM Assay. Scientific Reports 7, 10451, doi:10.1038/s41598-017-10856-3 (2017).

14 Sevrioukova, I. F. Apoptosis-Inducing Factor: Structure, Function, and Redox Regulation. Antioxidants & Redox Signaling 14, 2545–2579, doi:10.1089/ars.2010.3445 (2010).

15 Li, L. Z., Xu, H. N., Ranji, M., Nioka, S. & Chance, B. Mitochondrial redox imaging for cancer diagnostic and therapeutic studies. Journal of innovative optical health sciences 2, 325–341 (2009).

16 Wallrabe, H. et al. Segmented cell analyses to measure redox states of autofluorescent NAD(P)H, FAD & Trp in cancer cells by FLIM. Scientific Reports 8, 79, doi:10.1038/s41598-017-18634-x (2018).

17 Jiang, H. et al. Drug-induced oxidative stress in cancer treatments: Angel or devil? Redox Biology 63, 102754, 10.1016/j.redox.2023.102754 (2023).

18 Rodrigues, D. et al. Unravelling Mechanisms of Doxorubicin-Induced Toxicity in 3D Human Intestinal Organoids. International Journal of Molecular Sciences 23, 1286 (2022).

19 Durr, N., Ben-Yakar, A., Weisspfennig, C. & Holfeld, B. Maximum imaging depth of two-photon autofluorescence microscopy in epithelial tissues. Journal of Biomedical Optics 16, 026008 (2011).

20 Favreau, P. F. et al. Label-free redox imaging of patient-derived organoids using selective plane illumination microscopy. Biomed. Opt. Express 11, 2591–2606, doi:10.1364/BOE.389164 (2020).

21 Shirmanova, M. V. et al. Insight into redox regulation of apoptosis in cancer cells with multiparametric live-cell microscopy. Scientific Reports 12, 4476, doi:10.1038/s41598-022-08509-1 (2022).

22 Wang, L. et al. Doxorubicin-Induced Systemic Inflammation Is Driven by Upregulation of Toll-Like Receptor TLR4 and Endotoxin Leakage. Cancer Research 76, 6631–6642, doi:10.1158/0008-5472.Can-15-3034 (2016).

23 Kelly, C. Use of murine enteroids and colonoids to model gastrointestinal toxicity in response to anti-cancer agents in the small intestine and colon, University of Liverpool, (2024).

24 Zhao, Z. et al. Organoids. Nature Reviews Methods Primers 2, 94, doi:10.1038/s43586-022-00174-y (2022).

25 Nikolaev, M. et al. Homeostatic mini-intestines through scaffold-guided organoid morphogenesis. Nature 585, 574–578, doi:10.1038/s41586-020-2724-8 (2020).

26 Ambrosini, Y. M. et al. Recapitulation of the accessible interface of biopsy-derived canine intestinal organoids to study epithelial-luminal interactions. PLOS ONE 15, e0231423, doi:10.1371/journal.pone.0231423 (2020).

27 Mitrofanova, O. et al. Bioengineered human colon organoids with in vivo-like cellular complexity and function. Cell Stem Cell 31, 1175–1186.e1177, doi:10.1016/j.stem.2024.05.007 (2024).

28 Gonatas, N. K. & Avrameas, S. Detection of plasma membrane carbohydrates with lectin peroxidase conjugates. Journal of Cell Biology 59, 436–443, doi:10.1083/jcb.59.2.436 (1973).

29 Matute, J. D. et al. Intelectin-1 binds and alters the localization of the mucus barrier– modifying bacterium Akkermansia muciniphila. Journal of Experimental Medicine 220, doi:10.1084/jem.20211938 (2022).

30 Datta, S. S., Preska Steinberg, A. & Ismagilov, R. F. Polymers in the gut compress the colonic mucus hydrogel. Proceedings of the National Academy of Sciences 113, 7041–7046, doi:doi:10.1073/pnas.1602789113 (2016).

31 Dolan, B., Ermund, A., Martinez-Abad, B., Johansson, M. E. V. & Hansson, G. C. Clearance of small intestinal crypts involves goblet cell mucus secretion by intracellular granule rupture and enterocyte ion transport. Science Signaling 15, eabl5848, doi:doi:10.1126/scisignal.abl5848 (2022).

32 Koestler, B. J. et al. Human Intestinal Enteroids as a Model System of *Shigella* Pathogenesis. Infection and Immunity 87, 10.1128/iai.00733-00718, doi:doi:10.1128/iai.00733-18 (2019).

33 Gomez-Sjoberg, R., Leyrat, A. A., Houseman, B. T., Shokat, K. & Quake, S. R. Biocompatibility and Reduced Drug Absorption of Sol−Gel-Treated Poly(dimethyl siloxane) for Microfluidic Cell Culture Applications. Analytical Chemistry 82, 8954–8960, doi:10.1021/ac101870s (2010).

34 Michielin, F. et al. The Microfluidic Environment Reveals a Hidden Role of Self-Organizing Extracellular Matrix in Hepatic Commitment and Organoid Formation of hiPSCs. Cell Reports 33, doi:10.1016/j.celrep.2020.108453 (2020).

35 Löffler, J. et al. A comprehensive molecular profiling approach reveals metabolic alterations that steer bone tissue regeneration. Communications Biology 6, 327, doi:10.1038/s42003-023-04652-1 (2023).

36 Chandra, L. et al. Derivation of adult canine intestinal organoids for translational research in gastroenterology. BMC Biology 17, 33, doi:10.1186/s12915-019-0652-6 (2019).

37 Bulin, A.-L., Broekgaarden, M. & Hasan, T. Comprehensive high-throughput image analysis for therapeutic efficacy of architecturally complex heterotypic organoids. Scientific Reports 7, 16645, doi:10.1038/s41598-017-16622-9 (2017).

38 Martinez-Ordoñez, A. et al. Whole-mount staining of mouse colorectal cancer organoids and fibroblast-organoid co-cultures. STAR Protocols 4, 102243, 10.1016/j.xpro.2023.102243 (2023).

