## Supplementary materials for "OrganoidChip+: a Microfluidic Platform for Culturing, Staining, Immobilization, and High-Content Imaging of Adult Stem Cell-Derived Organoids"

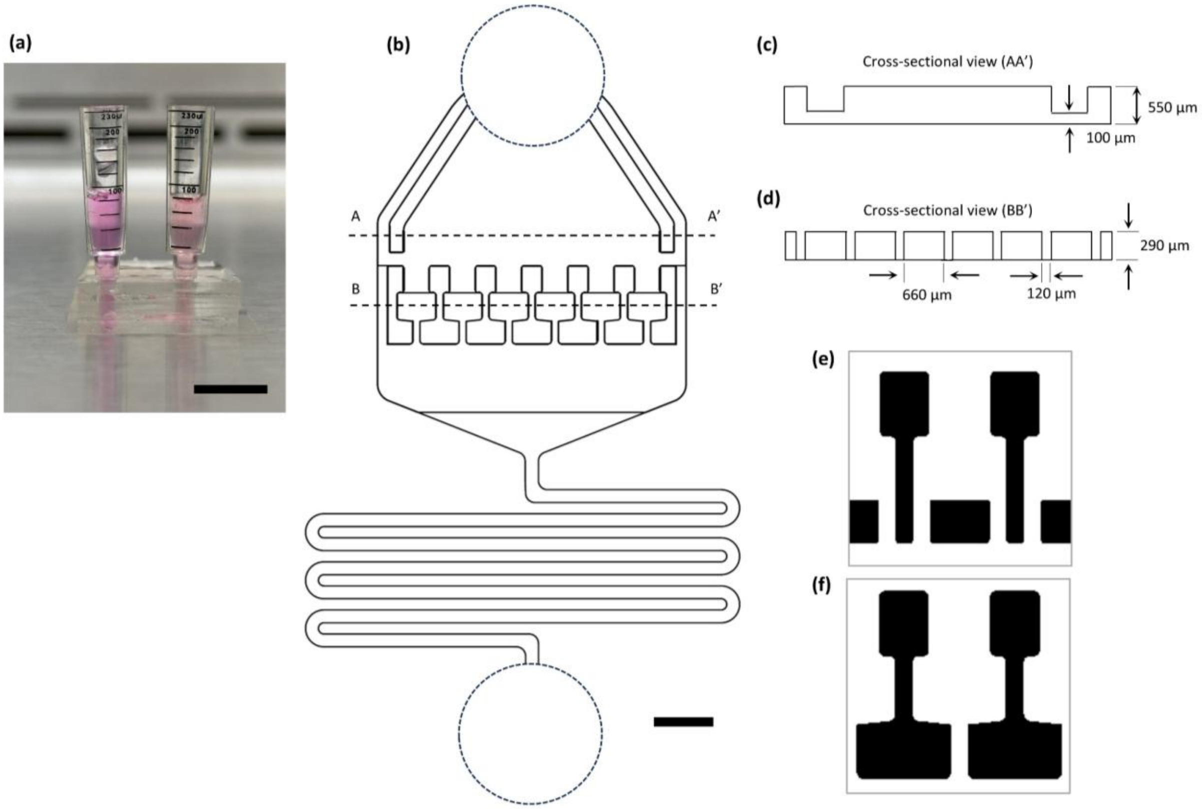


**Figure S1: (a)** The OrganoidChip+ with two 230 µl glass reservoirs mounted at the inlet and outlet that serve as the nutrient supply for the organoid culture. **(b)** depicts the secondary chip design with slight variation in the filter channel geometry. Cross sections AA' **(c)** and BB' **(d)** are shown with channel dimensions. The two filter channel designs from the primary **(e)** and secondary **(f)** chip design are shown. The PDMS microstructures are shown in black and channels in white. Scale bars in (a-b) represent 1 mm.

### Seeding density optimization

**Figure S2** depicts four experiments conducted in the original imaging OrganoidChip[1] that was designed without perfusion channels as tested with different seeding densities. The desired seeding densities were created by mixing the required number of clumps with Matrigel. We verified that the seeded densities closely followed the theoretical calculations obtained using a hemacytometer as described in the methods section (results not shown). The results indicated that at very high seeding densities (~ 448,000 clumps/mL of Matrigel, Exp 1), the growth rate of organoids was slower compared to those with lower seeding densities, preventing them from reaching sizes above 400 µm by Day 7 post-seeding. This reduced growth rate is likely due to higher nutrient demands of the larger number of cells, while nutrient supply remains limited by diffusion through Matrigel, thereby restricting organoid growth. By decreasing the seeding density (~ 385,000 clumps/mL of Matrigel, Exp. 2), the organoid sizes increase but the number of organoids exceeds the optimal level (8-12 organoids that are larger than 400 µm) for the chip. Conversely, at very low seeding densities (~ 25,000 clumps/mL of Matrigel, Exp. 4), organoids grow significantly larger than higher seeding densities, but the number of organoids is insufficient. A seeding density close to 150,000 clumps/mL (Exp. 2) generates a suitable number of organoids with desired sizes. Therefore, we chose to use this seeding density for subsequent experiments with the OrganoidChip+ presented in this study.

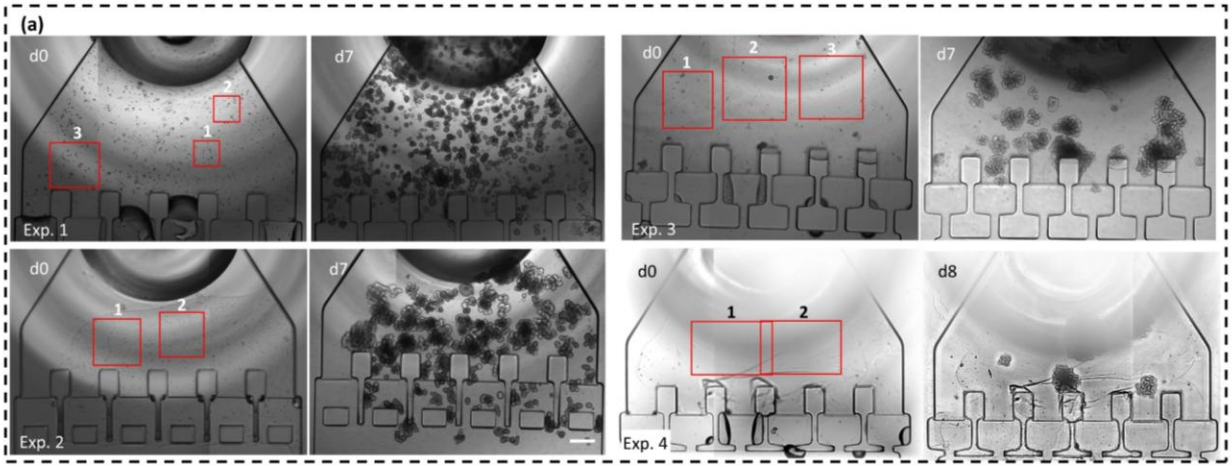


(b)

| Exp. # | 1 | | | 2 | | 3 | | | 4 | |
|---|---|---|---|---|---|---|---|---|---|---|
| ROI # | 1 | 2 | 3 | 1 | 2 | 1 | 2 | 3 | 1 | 2 |
| clumps/ml | 332,000 | 398,000 | 614,000 | 395,000 | 375,000 | 130,000 | 120,000 | 85,000 | 24,500 | 25,500 |
| Average clumps/ml | 448,000 | | | 385,000 | | 112,000 | | | 25,000 | |
| # of organoids with size > 400 µm | 0 | | | 13 | | 9 | | | 4 | |

**Figure S2: Organoid culturing with different initial seeding densities using the original imaging OrganoidChip designed without perfusion channels[1]. (a)** Four on-chip cultures of intestinal organoids on day 0 (left) and day 7 (right) with seeding densities decreasing from Exp. 1 to Exp. 4. The cell densities were calculated by directly measuring the number of cell clumps in each ROI, shown as red boxes, divided by the Matrigel volume within the ROI. **(b)** The cell clump densities in each ROI, the average seeding densities and number of organoids with size above 400 µm for each experiment. Scale bar in (a) indicates 400 µm.

### Organoid growth tracking and calculation

#### Image analysis and growth rate calculation

To track organoid size during growth, we captured brightfield images at different depths every day, obtained the minimum intensity projection image, and binarized this image to find the projected area of each organoid on each day (**Figure S3**). Also, each organoid was assigned a number after seeding. To compensate for the initial size of each organoid at seeding, we divide the projected areas of each organoid on each day to its area on Day 0, as shown in **Figure S3(d)** and defined in **Eq. S1**, below:

$$GR_i^n = \frac{A_i^n}{A_i^0} \qquad \text{S1}$$

Where GR is the growth rate, and "n" and "i" correspond to the number of days after seeding and organoid number, respectively. To find the mean growth rate in each off- or on-chip experiment at each day, we used the arithmetic mean of the growth rates as follows:

$$GR_{mean}^n = \frac{\sum_{i=1}^{m} GR_i^n}{m} \qquad \text{S2}$$

Where "m" is the total number of organoids in each experimental group. We also obtained the average growth rates across 7 off-chip and on-chip experiments according to **Eq. S3** below:

$$GR_{avg}^n = \frac{\sum_{k=1}^{7} GR_{mean,k}^n}{7} \qquad \text{S3}$$

where "k" is the experiment number. **Eqs. S1** to **S3** were calculated separately for off- and on-chip experimental groups.

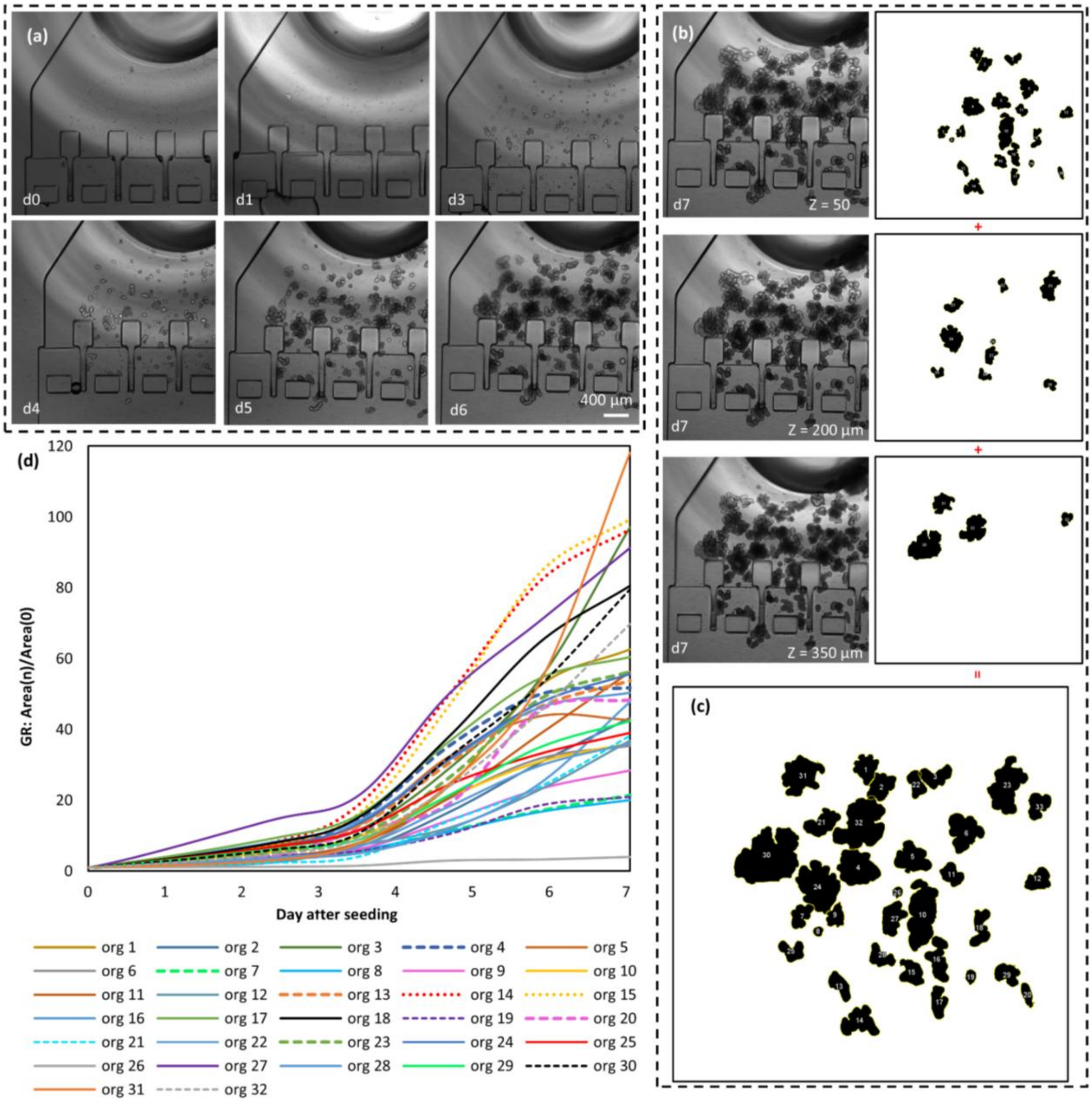


**Figure S3: Imaging and analysis of on-chip organoid growth. (a)** Time-lapse images of growth of organoids starting from cell clumps on day 0 into large mature organoids over the course of 6 days. **(b)** same FOV on day 7; images were captured at different depths to capture all in-focus organoids. The right column shows the binarized images of each corresponding brightfield image. **(c)** The binarized images in **(b)** were summed up to achieve **(c)**. Organoids were numbered (32 in total) to be able to track their growth over 7 days. **(d)** The areas of organoids were calculated from the binarized images. Then, the area on each day for each organoid was divided by their area at seeding for normalization. Area(n) is the area on the n$^{th}$ day.

### Heatmap calculations

For heatmap calculations, only growth rates at Day 7 were used. The growth rate for each organoid was normalized by the maximum growth rate on Day 7 among all organoids in that experimental group containing both off- and on-chip organoids grown in parallel, as defined in **Eq. S4**. This normalization preserves the relative growth rates while enabling consistent comparison across datasets with differing growth rate ranges.

$$GR_i^{norm} = \frac{GR_i^7}{Max\{GR_1^7,\ GR_2^7,\ \ GR_3^7, \dots\}} \qquad \text{S4}$$

The radius of the Matrigel domes in all experiments were averaged. A quarter of a circle with the averaged radius was used to create the off-chip heatmap. Due to the long analysis times required, we only analyzed a quarter of the Matrigel domes and projected all the analyzed organoids to the circle quarter with average radius for demonstration. The area of each organoid was used to calculate the radius of the circle with equal area. Therefore, each organoid was plotted as a circle (**Figs. 2e-f**). The heatmaps were generated by assigning the $GR_i^{norm}$ values and their respective colors to each organoid's corresponding circle in the heatmap matrix, $\mathrm{Z}(\mathrm{x},\mathrm{y})$. For areas in which multiple organoids were overlapped, the heatmap values were calculated by averaging the $GR_i^{norm}$ value between the overlapping organoids.

For on-chip heatmap generation, the same procedure used for off-chip heatmaps was applied, with the exception of incorporating plane symmetry. Since certain regions of the culture chamber lacked organoids in all 7 analyzed experiments, we leveraged the chip's plane symmetry to average growth rates across the two halves divided by the chip's centerline (**Fig. 2e**), enabling heatmap construction.

Next, we normalized the elements of each heatmap matrix by the maximum element value across both matrices (**Eq. S5**) to plot the heatmap color values between 0 and 1.

$$Z_{norm}(x, y) = \frac{Z(x, y)}{Max(Z_{max}^{on}\ ,\ Z_{max}^{off})} \qquad \text{S5}$$

where $\mathrm{Z}_{\mathrm{max}}$ is the maximum element value for the off- or on-chip heatmap matrix found. $\mathrm{Z}_{\mathrm{norm}}(\mathrm{x},\mathrm{y})$ is the final matrix generating the heatmap and is calculated for the off- and on-chip cases, separately.

For generating **Fig. 2g**, we first obtained the heatmap matrix for the off- and on-chip cases. For off-chip, the heatmap values were calculated along the radial dotted lines in **Fig. 2f**. The dotted lines start from the dome center and span radially to the dome boundary. Not all lines are displayed in **Fig. 2f**, as a total of 91 lines were used, with angles relative to the horizontal edge of the quarter ranging from 0˚ to 90˚ in 1-degree increments. The heatmap values along all the lines were assigned to matrix $A \in R^{k \times \omega}$, where k is the number of angles (=91) and ω is the number of equidistance points along radius whose values were obtained by interpolation. The non-zero elements in each column of matrix *A* were averaged to yield a 1 × ω vector. For on-chip, the heatmap values in half of the culture chamber were obtained and assigned to a matrix $B \in R^{\psi \times \upsilon}$ where $\psi$ and $\upsilon$ are the rows and columns shaping the culture chamber in the *y* and *x* directions, respectively. The non-zero elements in each row of matrix *B*, as they belonged to the culture chamber, were averaged to yield a $\psi \times 1$ vector. The two vectors were spanned over [0 R] and [0 Y] for the off- and on-chip cases, respectively, and plotted in **Fig. 2g**.

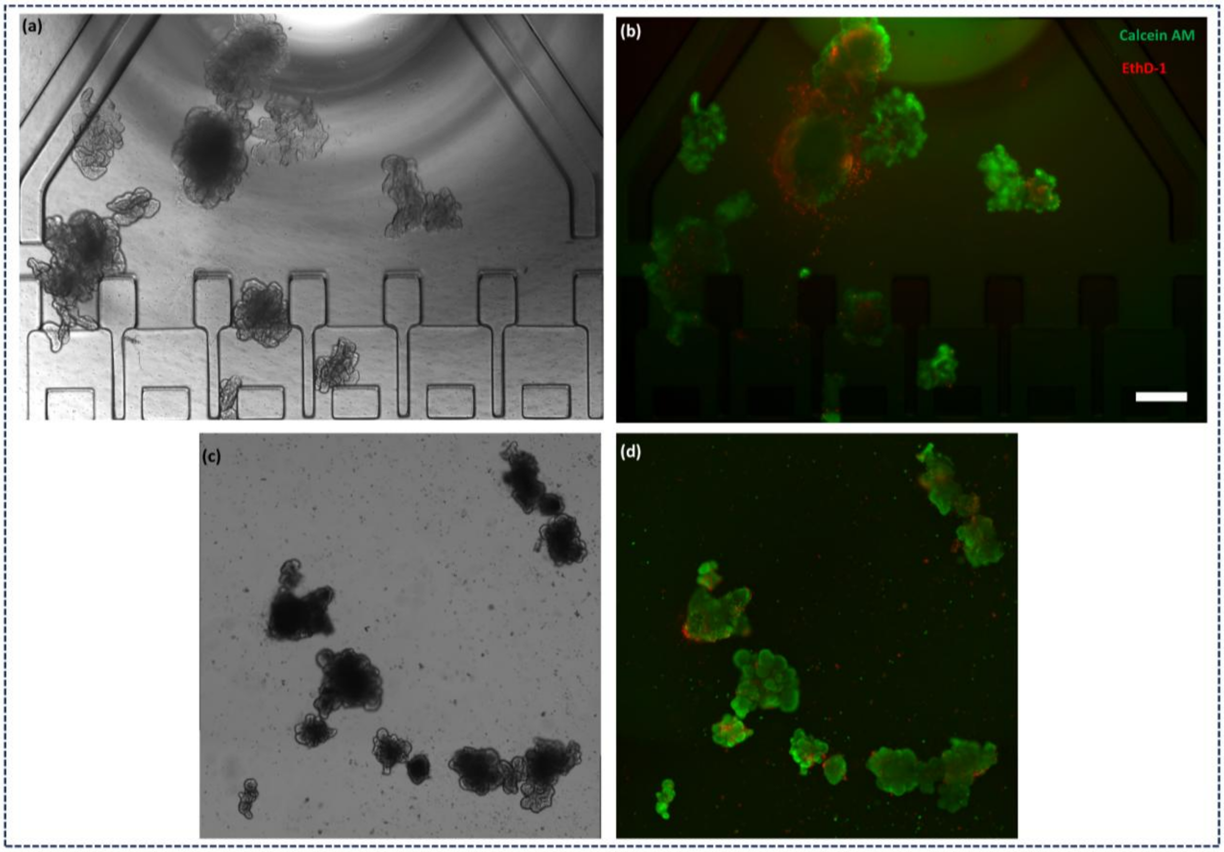


**Figure S4: Fluorescence viability assay on organoids grown on-chip and off-chip for 7 days. (a)** and **(c)** show brightfield images and figures **(b)** and **(d)** show fluorescence images of on- and off-chip organoids stained with Calcein AM and EthD-1 for live cell and dead cell staining, respectively. Scale bar in (b) represents 400 µm and applies to all subfigures.

### Two-photon image analysis and redox ratio calculations

To find the redox ratio values, we segmented the images captured using the 745 nm and 860 nm wavelengths to find the boundaries of the imaged organoid at each depth (z). For segmenting the organoid boundaries, we followed the flowchart in **Figure S5**. Next, we calculated the redox ratio for each segmented area by obtaining the mean pixel intensity within that area for each wavelength $\overline{S_{745}}$ and $\overline{S_{860}}$ (**S6**). Redox ratio calculation was performed according to **Eqs. S6-S12** as described in detail in the section below. The redox ratio at each depth was found by area-weighted averaging of the redox ratios of the segmented areas in that depth (**S6**). The redox ratios of all depths were averaged and presented as the redox ratio of that organoid. To find the redox ratio values plotted in **Figure 4c**, the redox ratio of the vehicle control organoids of the off- and on-chip groups were averaged (**S11**) and used in **S12** to normalize the redox ratio across each experiment. Three experiments were performed for each Dox concentration. All image processing and calculations were performed in Fiji and Microsoft Excel.

#### Detailed redox ratio calculations

Our in-house-built two-photon microscope is capable of reading one output at a time, therefore simultaneous imaging of NADH and FAD was not possible. Also, we wanted to avoid switching between emission filters for capturing NADH and FAD at each depth as that would unreasonably increase the imaging time. Therefore, we used a broadband bandpass filter (BG39, Schott) to collect both NADH and FAD signals. According to Qin *et al.*, the two-photon action cross-section of NADH at 860 nm is zero and the action potential of both molecules is relatively high at 745 nm[2]. Therefore, we chose excitation wavelengths of 745 nm, to excite both NADH and FAD, and 860 nm, to only excite FAD, and captured images at each wavelength at all depths. As seen in **S10**, we first normalized the pixel intensities by the square of the excitation powers. To decompose the intensity of NADH and FAD molecules, we had to carry out two scaling steps according to the action cross-section values to be able to compare pixel intensity values collected from the two wavelengths. For this purpose, we used two scaling factors, one for scaling the FAD signal from 860 nm to 745 nm (α, **S7**) and one for scaling the NADH signal to the FAD signal at 745 nm (β, **S8**). Next, we found the transmission power of the filter with respect to NADH and FAD emission spectra by calculating the area under the curve made by the product of the filter transmission and the emission curves (see **S9** and **Table S1**). The filter transmission curve was obtained from the manufacturer's website[3] and, the NADH and FAD emission spectra curves were found in the literature[4]. The quantum yield (q) of each molecule was estimated from Gorbunova et al. to be 2.1 and 2.6 for NADH and FAD, respectively[5].

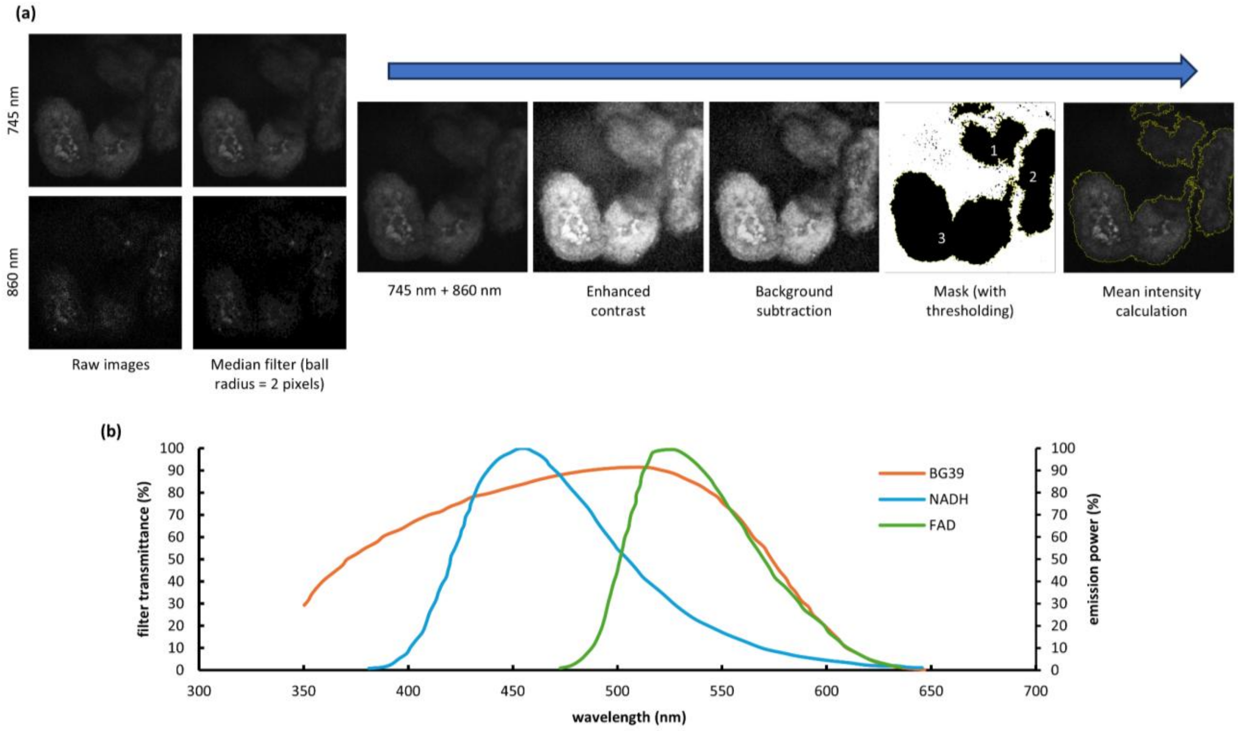


**Figure S5: (a)** Illustrates the image analysis for redox ratio calculations. First, the raw images are filtered with the median filter using a ball radius of 2 pixels. Next, the two images are summed, the contrast is enhanced, and the background is subtracted to find the area of the image occupied by the organoid(s). The resulting image is thresholded to mark the organoid boundaries and convert them to masks. The mask for each area is used to calculate the mean pixel intensity within that mask for images captured using the 745 nm and 860 nm excitation wavelengths. **(b)** shows the transmission efficiency of the BG39 filter (orange) along with the emission spectra of NADH (blue) and FAD (green). The values were digitized from the source plots*,[4] and an integration was performed according to S9 to find the amount of fluorescent light transmitted through the filter.

* https://www.edmundoptics.com/document/download/354140

$$\bar{S}(z) = \frac{\sum_i^n \sum_j^m S_{i,j}}{\sum_i^n A_i} \qquad \text{S6}$$

$$\alpha = \frac{CS_{745}^{FAD}}{CS_{860}^{FAD}} \qquad \text{S7}$$

$$\beta = \frac{CS_{745}^{FAD}}{CS_{745}^{NADH}} \qquad \text{S8}$$

$$\emptyset_l = \int_{w_1}^{w_2} E_l\, F\, dw\ , \text{"}l\text{" is } NADH \text{ or } FAD \qquad \text{S9}$$

$$RR = \frac{NADH}{FAD} = \frac{\left[\frac{\frac{\overline{S_{745}}}{P_{745}^2} - \frac{\overline{S_{860}}}{P_{860}^2}\alpha}{\emptyset_{NADH}\, q_{NADH}}\right]\beta}{\left[\frac{\frac{\overline{S_{860}}}{P_{860}^2}\alpha}{\emptyset_{FAD}\, q_{FAD}}\right]} \qquad \text{S10}$$

$$RR_{control}^{avg} = \frac{\sum_{j=1}^{n_{on}} RR_{vehicle}^{j} + \sum_{j=1}^{n_{off}} RR_{vehicle}^{j}}{n_{on} + n_{off}} \qquad \text{S11}$$

$$RR_{norm}^{i} = \frac{RR^i}{RR_{control}^{avg}} \qquad \text{S12}$$

See **Table S2** for abbreviations used in **Eqs. S6** to **S12**.

**Table S1: Integral bounds used for calculating $\phi$ in S9.**

| | $w_1$ (nm) | $w_2$ (nm) |
|---|---|---|
| NADH | 380 | 635 |
| FAD | 470 | 635 |

**Table S2: Shows the abbreviations used in equation S6 to S12 for the redox ratio analysis.**

| Abbreviation | description | Abbreviation | description |
|---|---|---|---|
| $S$ | pixel intensity | $P$ | laser power exposed on the sample |
| $\bar{S}(z)$ | average pixel intensity at a certain depth | $CS$ | action cross sections |
| $A$ | area | $E_I$ | emission intensity (a.u.) for NADH or FAD |
| $n$ | number of segmented regions | $F$ | filter transmission (%) |
| $m$ | number of pixels in each segmented region | $\phi$ | area under the curve of the product of the filter transmission and the emission curves |
| $RR$ | redox ratio | $\phi_{NADH}$ | $7.7 \times 10^5$ (a.u.) |
| $q$ | quantum yield | $\phi_{FAD}$ | $5.4 \times 10^5$ (a.u.) |
| $q_{NADH}$ | 2.1 | $\alpha$ | 2.57 |
| $q_{FAD}$ | 2.6 | $\beta$ | 3.70 |
| $RR_{vehicle}$ | Redox ratio of vehicle control groups | | |

**Table S3: Shows *p*-values of the *F*-tests performed on various redox ratio experimental groups for equivalence of the variances.**

| | | **on-chip** | | | **off-chip** | | |
|---|---|---|---|---|---|---|---|
| | | vehicle | 0.3 µM | 2 µM | vehicle | 0.3 µM | 2 µM |
| | vehicle | | 0.11 | 0.18 | 0.26 | | |
| **on-chip** | 0.3 µM | 0.11 | | 0.39 | | 0.34 | |
| | 2 µM | 0.18 | 0.39 | | | | 0.40 |
| | vehicle | 0.26 | | | | 0.40 | 0.22 |
| **off-chip** | 0.3 µM | | 0.34 | | 0.40 | | 0.35 |
| | 2 µM | | | 0.40 | 0.22 | 0.35 | |

**Table S4: Shows *p*-values of the *t*-tests performed on various redox ratio experimental groups for equivalence of the means.** *P*-values in red show unexpected test results.

| | | on-chip | | | off-chip | | |
|---|---|---|---|---|---|---|---|
| | | vehicle | 0.3 µM | 2 µM | vehicle | 0.3 µM | 2 µM |
| **on-chip** | vehicle | | 0.00297 | 0.00003 | 0.193 | | |
| | 0.3 µM | 0.00297 | | 0.00744 | | 0.208 | |
| | 2 µM | 0.00003 | 0.00744 | | | | 0.790 |
| **off-chip** | vehicle | 0.193 | | | | 0.00238 | 0.00005 |
| | 0.3 µM | | 0.208 | | 0.00238 | | 0.134 |
| | 2 µM | | | 0.790 | 0.00005 | 0.134 | |

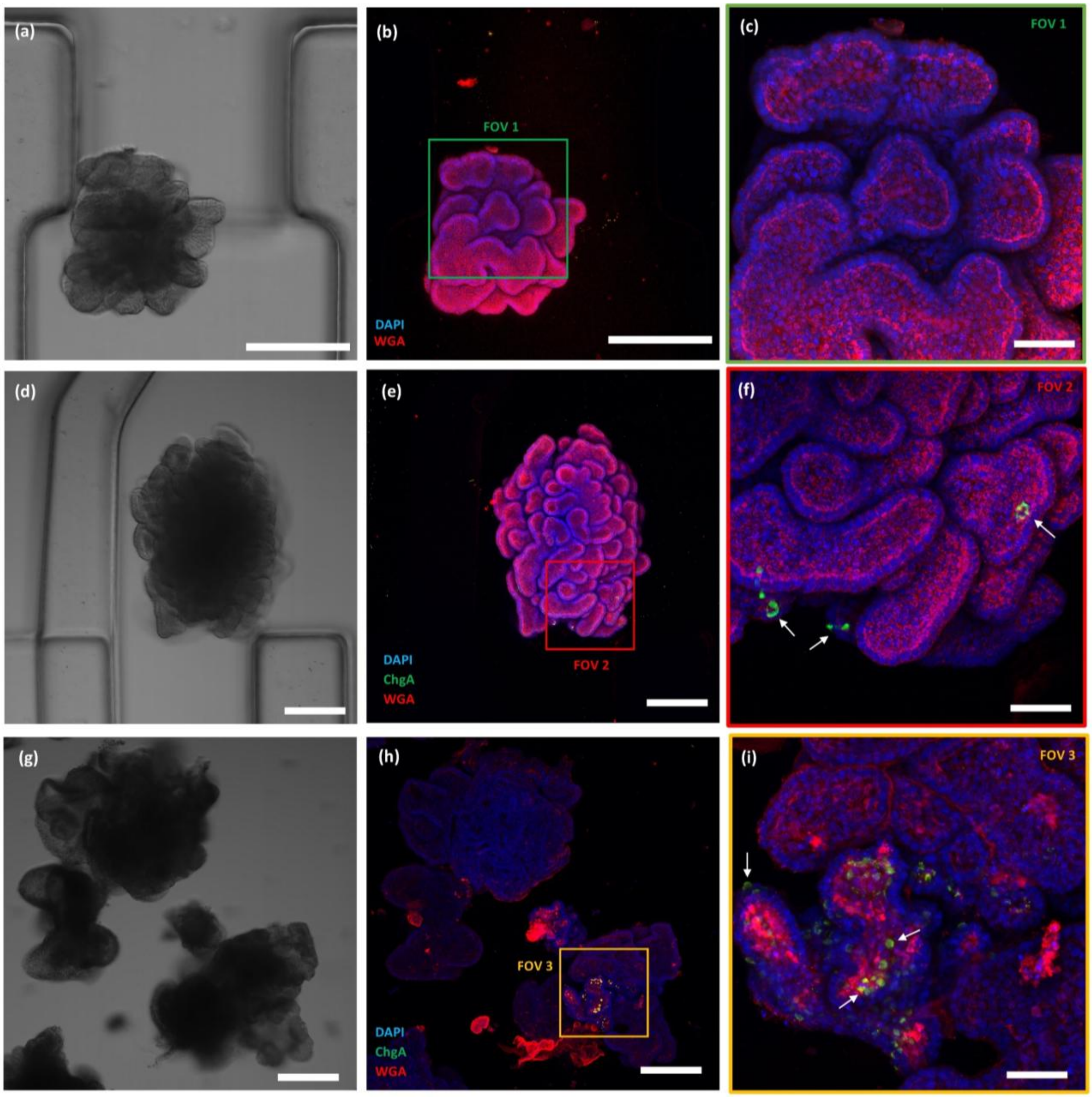


**Figure S6: Immunofluorescence, DAPI and WGA staining of organoids grown off- and on-chip. (a)** Shows the brightfield image of an organoid trapped in a TA and **(b)** shows the same organoid stained with DAPI and WGA (0.3 NA, 10×). **(c)** Depicts FOV 1 shown in **(b)** captured using a high-resolution objective (0.6 NA, 40×), **(d-f)** Brightfield and fluorescence confocal images of an organoid stained with DAPI, ChgA, and WGA on-chip. **(g-i)** Brightfield and fluorescence confocal images of an organoid stained with DAPI, ChgA, and WGA off-chip. White arrows point out the $ChgA^{+}$ enteroendocrine cells. Scale bars represent 200 µm in **(a-b)**, **(d-e)** and **(g-h)**, and 50 µm in **(c)**, **(f)** and **(i)**.

**Table S5: Tabulates the quantities of time, speed of spin coating, UV energy, and temperature used for the photolithography of the mold.**

| | SPIN COATING TIME AND SPEED | SOFT BAKE, TIME AND TEMPERATURE | EXPOSURE TIME AND ENERGY | POST EXPOSURE BAKE, TIME AND TEMPERATURE | DEVELOPING TIME |
|---|---|---|---|---|---|
| **LAYER 1: SU8-2100** | 5 s, 500 rpm then 33 s, 3075 rpm | 5 min, 65 °C then 20 min, 95 °C | 240 mJ/cm$^3$ | 5 min, 65 °C 10+5 min, 95 °C | none |
| **LAYER 2: SU8-2100** | 5 s, 500 rpm then 33 s, 2300 rpm | 5 min, 65 °C then 45 min, 95 °C | 315 mJ/cm$^3$ | 5 min, 65 °C 25 min, 95 °C | none |
| **LAYER 3: SU8-2100** | 5 s, 500 rpm then 33 s, 1850 rpm | 7 min, 65 °C then 56 min, 95 °C | 370 mJ/cm$^3$ | 5 min, 65 °C 20 min, 95 °C | 50 min (changing developer every 15 minutes) |

### References


1 Moshksayan, K. *et al.* OrganoidChip facilitates hydrogel-free immobilization for fast and blur-free imaging of organoids. *Scientific Reports* **13**, 11268, doi:10.1038/s41598-023-38212-8 (2023).

2 Qin, Y. & Xia, Y. Simultaneous Two-Photon Fluorescence Microscopy of NADH and FAD Using Pixel-to-Pixel Wavelength-Switching. *Frontiers in Physics* **9**, doi:10.3389/fphy.2021.642302 (2021).

3 AG, S. Coating Curve: BG-39 Colored Glass Bandpass Filter Internal Transmittance Graph. *Edmund Optics Inc.* (2009).

4 Huang, S., Heikal, A. A. & Webb, W. W. Two-Photon Fluorescence Spectroscopy and Microscopy of NAD(P)H and Flavoprotein. *Biophysical Journal* **82**, 2811-2825, doi:10.1016/S0006-3495(02)75621-X (2002).

5 Gorbunova, I. A. *et al.* Determination of quantum yield of NADH and FAD in alcohol-water solutions: the analysis of radiative and nonradiative relaxation pathways. *arXiv preprint arXiv:2205.00367* (2022).
